# Kinetochore life histories reveal the origins of chromosome mis-segregation and correction mechanisms

**DOI:** 10.1101/2021.03.30.436326

**Authors:** Onur Sen, Jonathan U. Harrison, Nigel J. Burroughs, Andrew D. McAinsh

## Abstract

Chromosome mis-segregation during mitosis leads to daughter cells with deviant karyotypes (aneuploidy) and an increased mutational burden through chromothripsis of mis-segregated chromosomes. The rate of mis-segregation and the aneuploidy state are hallmarks of cancer and linked to cancer genome evolution. Errors can manifest as “lagging chromosomes” in anaphase, although the mechanistic origins and likelihood of correction are incompletely understood. Here we combine lattice light sheet microscopy, endogenous protein labelling and computational analysis to define the life history of > 10^4^ kinetochores throughout metaphase and anaphase from over 200 cells. By defining the “laziness” of kinetochores in anaphase, we reveal that chromosomes are at a considerable and continual risk of mis-segregation. We show that the majority of kinetochores are corrected rapidly in early anaphase through an Aurora B dependent process. Moreover, quantitative analyses of the kinetochore life histories reveal a unique dynamic signature of metaphase kinetochore oscillations that forecasts their fate in the subsequent anaphase. We propose that in diploid human cells chromosome segregation is fundamentally error prone, with a new layer of early anaphase error correction required for stable karyotype propagation.

## Introduction

High-fidelity chromosome segregation is a key task during mitosis, and is crucial for maintaining the correct number of diploid chromosomes in a cell. Errors in chromosome segregation can lead to aneuploidy, the deviation in chromosome number from the diploid state. Such changes to the karyotype are associated with cancer progression, developmental disorders and ageing^1–3^. These pathologies are the consequence of gene dosage changes and proteomic stress^4,5^. Mis-segregated whole chromosomes can also give rise to micronuclei that are distinct from the main daughter cell nuclei. The micronucleus can be a site of further mutational processes due to aberrant replication of the physically isolated chromosome, leading to chromothripisis, which is characterised by extensive genomic rearrangements^6,7^. Like whole chromosome aneuploidy, chromothripisis is also associated with the evolution of human disease processes.

Avoiding aneuploidy requires that all replicated chromosomes (sister chromatids) are accurately segregated into two daughter cells during mitosis. High-fidelity segregation is founded on two key principles, 1) Back-to-back geometry of sister kinetochores (multi-protein machines assembling on each sister centromere) favor binding to microtubules emanating from opposite spindle poles (amphitelic attachments)^8^, 2) Error correction mechanisms detect and destabilise improper attachments^9–11^ and the spindle assembly checkpoint (SAC) delays anaphase onset until correct attachments are achieved^12^. A key challenge is to understand how and why some erroneous kinetochore-microtubule attachments can escape these surveillance mechanisms.

It is well established that merotelic attachment is a major source of aneuploidy^13^, in which one, or both, sister kinetochores form attachments to both spindle poles during prometaphase^14^. This can even cause stretching of the merotelically attached kinetochore due to pulling forces toward opposite poles^13,15^. The resulting microtubule occupancy and tension can also satisfy the spindle assembly checkpoint (SAC)^13^. As a result, cells initiate anaphase with these merotelic kinetochores appearing to “lag” behind the segregating clusters of poleward moving kinetochores^3^,16. It is these lagging chromosomes that are at high risk of forming micronuclei as the nuclear envelope reassembles^6^.

Why do merotelic attachments form? In prometaphase, as kinetochores undergo search-and-capture there is a probability of binding microtubules emanating from opposite poles^17^. The spindle geometry at nuclear envelope breakdown has been shown to affect this probability with reduced distance between spindle poles increasing the fraction of improper attachments^18–20^. There is also evidence that the rate of microtubule-kinetochore turnover is important with a balance between having the necessary stability to enable chromosome movement, and sufficient turnover to limit the lifetime of improper attachments^21^. The turnover rate is cell type specific and provides one explanation for increased chromosomal instability in cancer cells^22^. Physical properties of the kinetochore and the chromosome arms are also important, with increasing size elevating the risk of merotely and mis-segregation, respectively^23,24^. Importantly, errors are not limited to merotely. For example, prolonged metaphase delay can lead to premature sister chromatid separation (PSCS) due to cohesion fatigue^25,26^ and non-resolved syntelic attachments have been proposed as a source of non-disjunction^27–29^. However, the systematic detection of these different error events in pre-anaphase cells and establishing the causal relationships with segregation behavior in subsequent anaphase remains unresolved.

Error correction mechanisms provide a route to detect and eliminate improper microtubule attachments. Aurora B kinase, as a component of the chromosome passenger complex (CPC), localizes to the centromere-kinetochore interface^30^, and plays a central role in error correction during early mitosis^31^. If Aurora B activity is compromised, the frequency of syntelic and merotelic attachments increase due to the failure to correct improper attachments^32–35^. Aurora B is preferentially enriched on aberrantly attached kinetochores^36^ where it phosphorylates outer kinetochore proteins, to destabilise these erroneous attachments^37,38^. As sister kinetochores form amphitelic attachments, increased tension and intersister distance have been proposed to reverse the microtubule destabilising phosphorylations^39^, and ultimately lead to attachment stabilisation, which gradually increases during metaphase^40–44^. At the metaphase to anaphase transition, the motor protein Mklp2 relocates Aurora B to the spindle midzone^45^, where it generates a phosphorylation gradient^46^. This phospho-gradient has been shown to delay chromosome decondensation and nuclear envelope reassembly (NER)^47^. Alongside the potential for resolution of merotely based on imbalanced pulling forces^16^, there are clearly dedicated mechanisms to limit the risk of mis-segregation.

The current paradigm in the field is that only rare kinetochores can escape pre-anaphase surveillance in near-diploid human cells (RPE1, HCT116) with the resulting lagging chromosome rate of 5-7%, previously shown by fixed cell imaging^22,24,28^. However, how lagging chromosomes are defined is imprecise and often reliant on endpoint assays using fixed cell imaging, which do not detect chromosomes that lag after, or are corrected before, the fixation. Here, we use a combination of lattice light sheet imaging and computational analysis which enable a novel quantitative measure of chromosome lag to be defined. We term this “laziness” and reveal how a much larger proportion of kinetochores are at risk of mis-segregation than previously thought. We also show that Aurora B operates during early anaphase to promote the rapid correction of the majority of lazy kinetochores. By analysing the history of lazy kinetochores, we identify key dynamic signatures in metaphase that predict the ultimate segregation outcome. These data provide new insight into the origins of chromosome mis-segregation in diploid human cells.

## Results

### Lattice light sheet imaging and automated analysis tools allow us to probe the origins of chromosome segregation errors

To understand the origins of chromosome segregation errors during mitosis, we used lattice light sheet microscopy^48^ to collect full 3D volumes every 4.7 seconds (s) for a total duration of between 3.1 and 17.2 minutes (mins) (median 8.3 mins). These image sequences capture events from late prometaphase through metaphase and anaphase onset to the late stages of anaphase (Fig. 1a; Movie S1). For this, we used a non-transformed near-diploid human hTERT-RPE1 cell line in which one allele of the NDC80 gene is tagged with eGFP at the carboxy-terminus^49^. By adapting our existing kinetochore tracking (KiT) algorithms^50^, we were able to capture an average of 34±9 long tracks of paired sister kinetochores per cell, compared to the total diploid number (46 pairs), representing 74 ± 20%, (Fig. 1b, Supp. Fig. 1). These long tracks lasted at least 75% of the duration of each movie providing an unprecedented near-complete 3D view of kinetochore dynamics. All of the imaged cells entered anaphase, demonstrating that there were no phototoxicity effects in our imaging setup (see methods for details). Furthermore, population level analysis of kinetochore trajectories in metaphase confirms that sister kinetochore pairs underwent heterogenous oscillatory motion with a half period of ∼40 s as previously described (Fig. 1c, 1d and 1e, Supp. Fig. 2a)^51,52^. This lattice light sheet imaging and analysis pipeline thus capture dynamics of kinetochores at high temporal resolution from late prometaphase to late anaphase.

**Figure 1.**
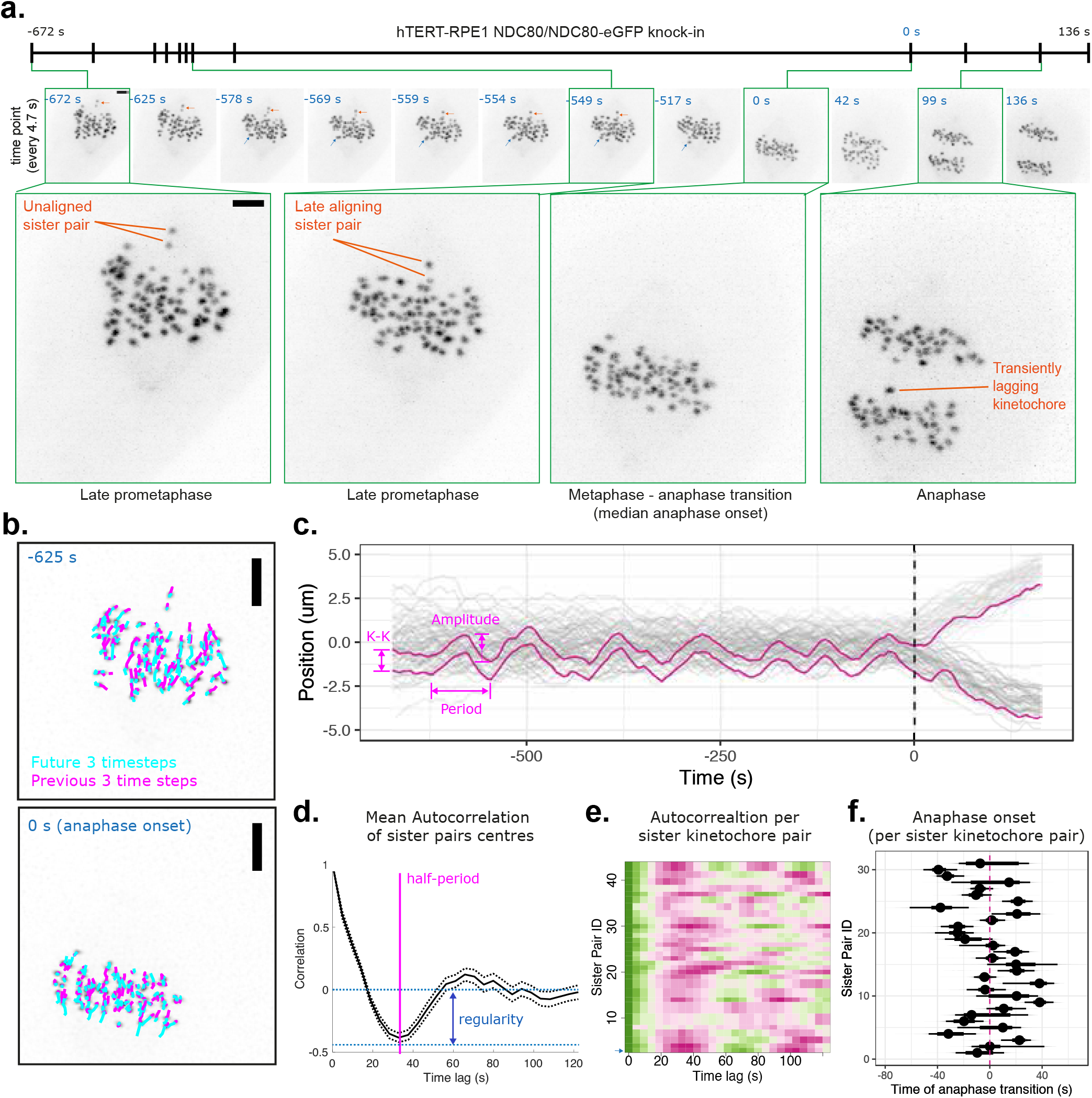
Lattice light-sheet microscopy enables imaging of mitosis with high spatiotemporal resolution and quantitative analysis of kinetochore dynamics. **a**, (top panel) *Z*-projected movie stills of a cell progressing from late prometaphase to anaphase (time 0 = median anaphase onset time for this cell). Scale bar is 2 µm. Time resolution is 4.7 s/frame. Blue arrow annotates a kinetochore pair while it is being bi-oriented. Red arrow annotates a late aligning kinetochore pair. (bottom panel) Zoomed images from selected time points showing a late aligning sister pair and a transiently lagging kinetochore. Figures 1a-f show images and plots from the same cell. Scale bar is 3 µm. **b**, Example of kinetochore tracking using dragontails to annotate the previous and future 3 timesteps in *Z*-projected movie stills. Scale bar is 3 µm. **c**, Track overlay (*x*-axis position in time) shows trajectories of all sister kinetochore pairs (sister one in dark gray; sister two in light gray) during metaphase and anaphase. Magenta trajectories show the positions of a representative sister kinetochore pair that exhibit dynamic oscillations and timely segregation. Dashed line (time = 0) annotates the median anaphase onset time for this cell. Parameters related to kinetochore oscillations, intersister (K-K) distance, amplitude and period, are shown. **d**, Graph shows mean autocorrelation for all sister pair centres (kinetochore oscillations) in this cell. The time when the the autocorrelation curve achieves its minimum (a negative valley) corresponds to the half period of average kinetochore oscillations in this cell (35-40s, annotated with pink line), and the depth at the minimum is the oscillation regularity (blue arrow). **e**, Heat-map demonstrates the autocorrelation values (high values in green; low values in magenta) for each sister pair (each individual row) in this cell. Heat-map for the autocorrelation of the representative sister pair highlighted in (c) is plotted as PairID1 (bottom row), and annotated with blue arrow. **f**, Median and 95% credible intervals are shown for the anaphase onset time of each sister pair. Dashed line indicates the median anaphase onset time for this cell.

**Figure 2.**
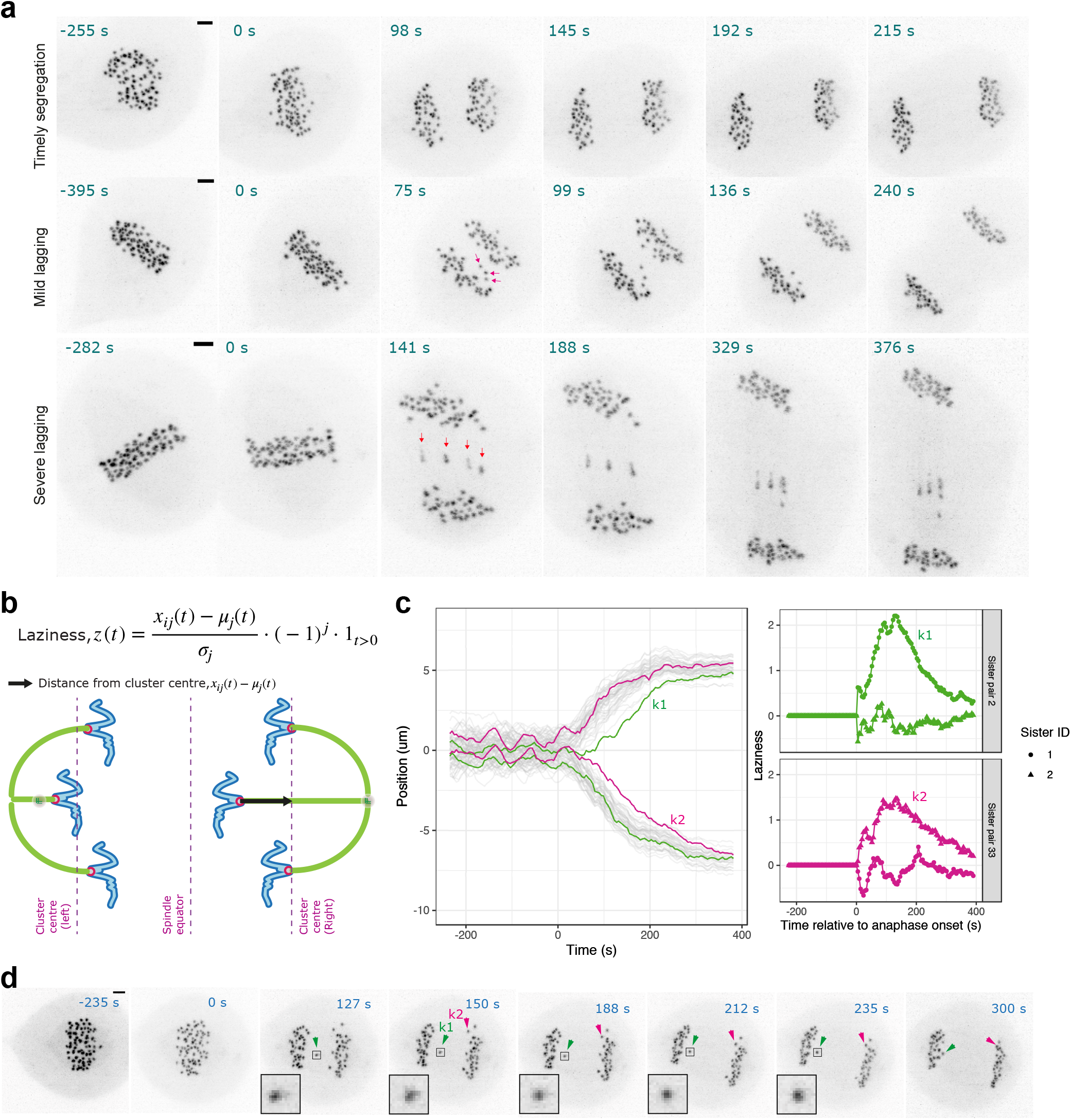
Laziness is a novel quantitative measure for spatiotemporal analysis of lagging chromosome behavior. **a**, *Z*-projected movie stills of representative cells that undergo timely segregation or segregation with mild or persistent lagging chromosomes. (middle panel) Magenta arrows annotate three kinetochores with different levels of lagging that are later corrected in anaphase. (bottom panel) Red arrows annotate multiple persistent lagging kinetochores that are not corrected in anaphase. **b**, Formula used for calculating laziness (z) of a lagging kinetochore is illustrated on a cartoon (see methods for details). **c**, (left panel) Track overlay (*x*-axis position in time) of a representative cell showing two highlighted sister pair trajectories annotated for kinetochores k1 and k2. (right panel) Laziness trajectory for k1 and k2 kinetochores and their sisters. **d**, *Z*-projected movie stills of the cell given in (c). Green arrowhead annotates kinetochore k1 and magenta arrowhead annotates kinetochore k2. Scale bar is 2 µm.

Identifying the causal events involved in chromosome mis-segregation is hampered by the low rate of chromosome mis-segregation in hTERT-RPE1 cells^24^. To circumvent this, we used a standard nocodazole arrest-and-release procedure in order to increase improper attachments. We also confirmed that the nocodazole arrest-and-release had little impact on oscillation dynamics in metaphase (Supp. Fig. 2a-f) suggesting kinetochore function is largely preserved.

To allow comparison of chromosome segregation behaviour between cells, we developed an algorithm that segments trajectories into metaphase and anaphase. Estimates of anaphase onset times for the kinetochore pairs are shown in Fig. 1f for the example cell displayed in Fig. 1a. The asynchrony in anaphase onset times between different sister pairs, as assessed by the median absolute deviation (a measure of spread similar to standard deviation but robust to outliers), was 16±6 s for all cells (*N* = 153 cells; DMSO, 2h noc, 4h noc pooled), consistent with earlier observations^51^. Lagging chromosomes then manifest as a kinetochore that is delayed in segregating, or fails to segregate at all, compared to the two clusters of kinetochores that are moving towards opposite spindle poles (e.g. Fig. 2a). Again, it is evident that defining kinetochores as “lagging” can be imprecise, i.e. when does late segregation become lagging? Indeed, we observed that kinetochores can exhibit a range of behaviors from timely segregation, and mild through to persistently lagging (Fig. 2a). These different levels of “chromosome lag” may not be consistently identified in manual assessments by different experts in absence of a quantifiable definition.

### A novel quantitative measure for spatio-temporal analysis of lagging chromosome behavior

To study factors that affect the fidelity of chromosome segregation systematically and without bias, we developed an automated tool to identify lagging chromosomes from time-series data. We assigned a quantitative measure, termed “laziness”, to reflect the individual segregation behavior of a kinetochore throughout anaphase. This allows us to analyse individual trajectories of kinetochores, and in this way reveal the time-evolution of laziness during anaphase. We define the laziness, *z*, for an individual kinetochore at any given time-point, based on its distance from the centre of the cluster of kinetochores to which it belongs; a cluster comprises the kinetochores destined for one of the daughter cells (Fig. 2b and methods). As a kinetochore’s lag increases behind its poleward moving cluster, the frame-to-frame laziness for that kinetochore takes increasingly high values (see the example lagging kinetochore *k*1 in Fig. 2c; Movie S2). This laziness over time is consistent with the movie image sequence of the same cell shown in Fig. 2d. For most lagging kinetochores, laziness increases to a peak value, and subsequently reduces over time as the kinetochore returns to its segregating cluster later in anaphase (e.g. *k*1 in Fig. 2c,d - see investigation of correction mechanisms below). On the other hand, other kinetochores can have similar laziness dynamics except that the maximum value reached is lower (compare *k*2 with max. laziness = 1.5 vs. *k*1 with max. laziness = 2.2, in Fig. 2c) and are not obviously lagging from the movie image sequence (*k*2 in Fig. 2d). The laziness thus quantifies anaphase chromosome behavior, and provides firm ground to investigate the underlying mechanisms.

We next calculated the maximum laziness for each kinetochore throughout anaphase. This provides a read-out that is independent of the duration and timing of a laziness event. The distribution of maximum laziness over the population of all kinetochore trajectories shows a sharp drop off and a long tail (*n* = 15576; *N* = 153 cells; Fig. 3a). As expected, we found a higher proportion of kinetochores with high maximum laziness after nocodazole arrest-and-release (pink lines). While there is aclear spectrum of laziness, we investigated whether there is also a distinct population of kinetochores that exhibit high laziness. For this, we fitted the maximum laziness scores to a generalized extremal value distribution. Based on quantile-quantile (Q-Q) plots, we find that beyond a laziness threshold, *a*, of approximately 2, the quality of the fit breaks down (Supp. Fig. 3). This suggests that there is a distinct population of kinetochores that reach higher laziness during anaphase.

**Figure 3.**
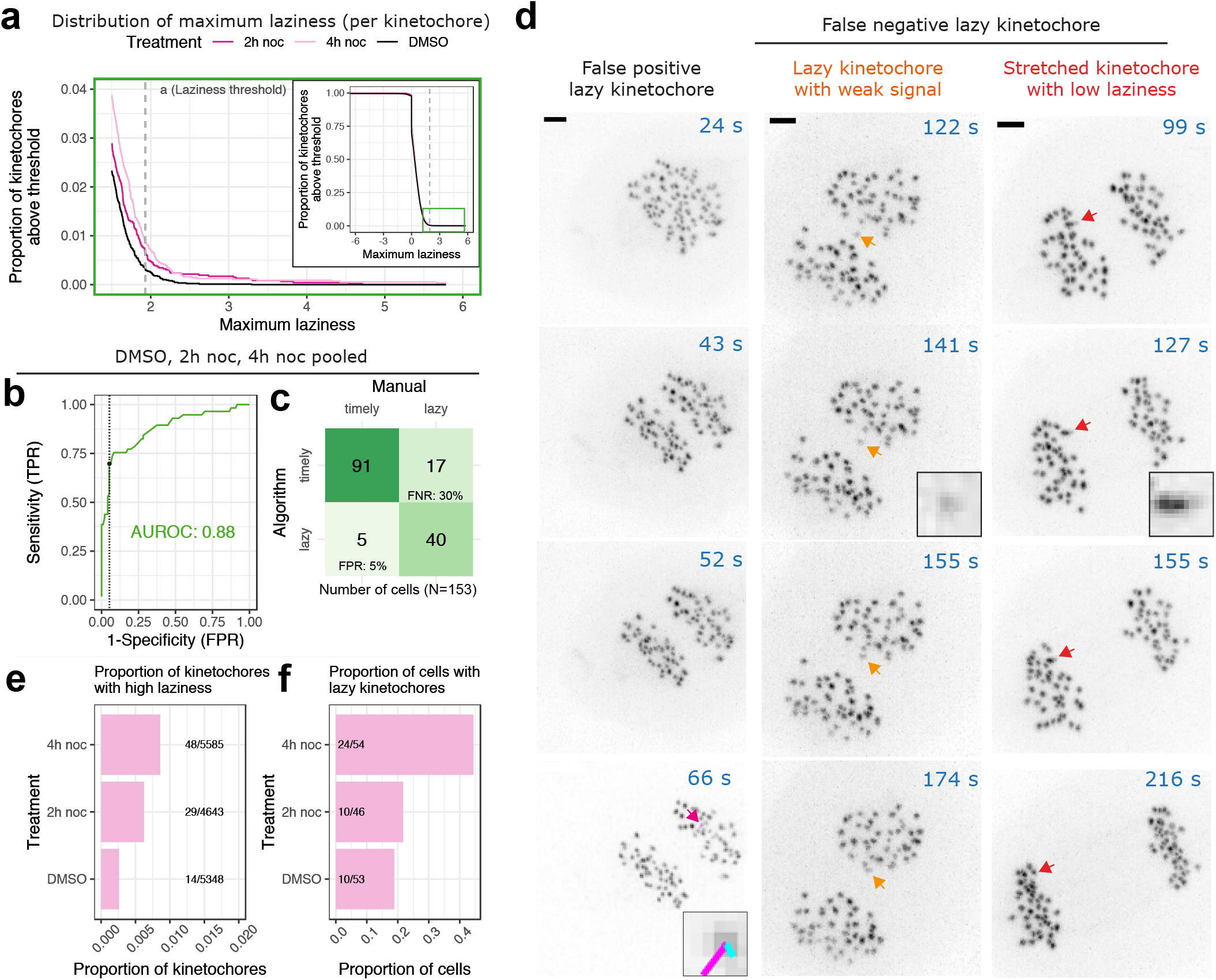
Calibration of the laziness threshold to manual assessment of chromosome segregation. **a**, Graph shows the distribution of maximum laziness (from laziness=1.0 to 6.0) reached by each kinetochore throughout anaphase, over the population of all kinetochore trajectories from three treatment groups; DMSO, 2h noc, 4h noc. Dashed line denotes laziness threshold. (top-right corner) Graph shows the distribution of all maximum laziness values. **b**, Graph shows receiver operator characteristic (ROC) curve at various laziness threshold settings. The laziness threshold (*a* = 1.93) is denoted by the black dot, and the corresponding false positive rate (5%) is denoted by the dashed line. The area under the ROC curve (AUROC) indicates how good the model is at distinguishing between the cells with at least one lazy kinetochore (reached laziness > *a* during anaphase) and the cells without lazy kinetochores, (timely). **c**, Confusion matrix demonstrates the number of cells (*N* = 153) having at least one lazy kinetochore (lazy), or zero lazy kinetochores (timely) during anaphase, quantified by the algorithm or manual assessment. False positive rate (FPR) indicates the cells scored as (having) lazy (kinetochore) by the algorithm but not by manual assessment. False negative rate (FNR) indicates the cells scored as lazy (having lazy kinetochore(s)) by manual assessment but not by the algorithm. **d**, *Z*-projected movie stills of representative cells that (left panel) have a false positive lazy kinetochore annotated with magenta arrow and magenta-cyan dragontails indicated by the algorithm; (middle panel) false negative lazy kinetochore, not detected by the algorithm due to low signal-to-noise ratio (zoomed kinetochore annotated by orange arrow); (right panel) false negative lazy kinetochore, scored as lazy by manual assessment due to merotely indicated by kinetochore stretching, but not by the algorithm due to being close to the cluster centre (zoomed kinetochore annotated by red arrow). **e**, Graph shows the proportion of lazy kinetochores (with max laziness score > *a*) in the cells treated with DMSO, 2h nocodazole or 4h nocodazole prior to washout. Numbers of lazy kinetochores and total kinetochores are shown for each treatment group. **f**, Graph shows the proportion of cells, that have at least one lazy kinetochore, treated with DMSO, 2h nocodazole or 4h nocodazole prior to washout. Numbers of cells with lazy kinetochore(s) and total cells are shown for each treatment group.

To further refine our estimate of this laziness threshold, *a*, and assess performance of the algorithm in identifying lagging kinetochores, we compared the output to (expert) manual inspection (Fig. 3b). For this, we plotted a receiver operator characteristic (ROC) curve at various threshold settings (of *a*). The area under this curve (AUC) is a performance indicator that quantifies how good the performance of the algorithm is against manual inspection at distinguishing between cells with lagging kinetochores and those without. Random chance would achieve an AUC of 0.5, whereas using the maximum laziness gives an AUC of 0.88 (Fig. 3b). Moreover, we can use the ROC curve to select a laziness threshold that gives us a false positive rate (FPR, segregation delays detected by the algorithm, but not by the manual inspection; for example Fig. 3d) of only ∼5% (Fig. 3c). This threshold (*a* = 1.93) is close to the approximate value (∼2) derived from the extremal distribution (Fig. 3a, Supp. Fig. 3).

With this threshold, we incur a false negative rate of ∼30% (FNR, scored as lagging by manual assessment but not by the algorithm) although visual inspection showed that these events are largely missed due to mis-tracking as a result of low signal-to-noise ratio, or due to manual scoring based on kinetochore stretching (merotely) rather than solely on the distance of a kinetochore to the cluster (e.g. in Fig. 3d). While this false negative rate reduces the number of segregation errors scored by the algorithm, we prioritize minimizing false positives which could distort downstream analysis.

This threshold, *a*, allows kinetochores to be classified either as lazy, (laziness > *a*) or timely (laziness < *a*). Lazy kinetochores thus “lag” a significant distance behind their segregating cluster at some point during anaphase. This analysis reveals that the proportion of lazy kinetochores in DMSO-treated cells is 0.0026, which increases (3.3 fold) to 0.0086 in nocodazole-treated cells in a duration dependent manner (Fig. 3e). We also quantified the proportion of cells that contain at least one lazy kinetochore at any point during anaphase, which is 0.18 for DMSO-treated cells, and increases to 0.44 with nocodazole treatment (Fig. 3f). We note this is higher than the proportion of untreated cells with lagging chromosomes (0.05-0.07) found by fixed cell imaging in previous reports^22,24,28^. Thus, the number of chromosomes at risk of mis-segregation in human cells is considerably higher than previously thought.

### Lazy kinetochores have a distinct dynamic mitotic signature

Quantitative analysis of lattice light sheet imaging provides 3D trajectory data going back in time to late prometaphase/metaphase. This opens up the possibility of identifying if there are any dynamic signatures in pre-anaphase kinetochores that can explain why certain kinetochores subsequently lag during anaphase. To do this, we separated kinetochores into those that underwent timely segregation (with maximum laziness ≤*a*) and those that exhibited segregation errors (with maximum laziness > *a*), henceforth referred to as lazy kinetochores. The kinetochore trajectory segments corresponding to metaphase were then extracted and analyzed. During metaphase, sister kinetochores undergo quasi-periodic oscillations along the spindle axis (Supp. Fig. 2a, b) with the distance between the two sisters also breathing (Supp. Fig. 2c, d) as kinetochores come under varying pulling and pushing forces^53–55^. We found that lazy kinetochores display reduced intersister (K-K) distance during metaphase which persists through the metaphase-anaphase transition (Fig. 4a, b, c). Moreover, the lazy kinetochore population also initiated anaphase (sister separation) 14 s later (relative to the average time for kinetochore pairs in that cell), was located on average closer to the metaphase plate, and moved poleward with a lower speed in anaphase. In contrast, there was no significant difference in the oscillation amplitude, speed, position in the metaphase plate (radius) or twist (angle between inter-kinetochore axis and normal to metaphase plate) (Fig. 4a). However, oscillations of lazy kinetochores were perturbed with a reduction in the regularity (Fig. 4d, compare Fig. 1d).

**Figure 4.**
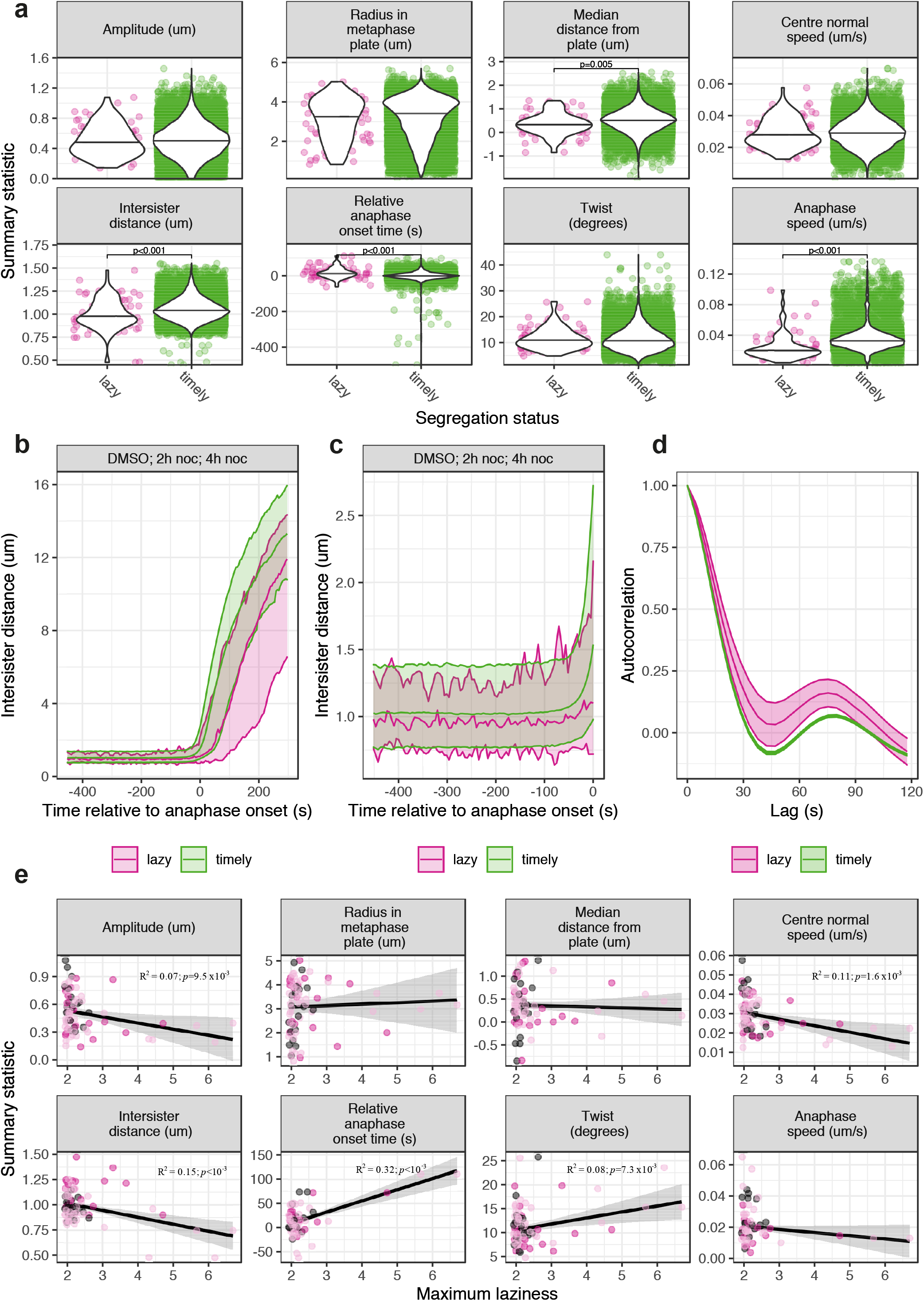
Lazy kinetochores have a distinct dynamic metaphase signature. **a**, Violin plots show medians of eight metaphase-anaphase variables (summary statistics) in lazy (with > *a* maximum laziness) and timely (with < *a* maximum laziness) kinetochores pooled from cells treated with DMSO, 2h noc or 4h noc. **b**, Graph shows median intersister (K-K) distance (bands show 2.5%, 50% and 97.5% quantiles) of all timely and lazy kinetochores throughout metaphase and anaphase. **c**, Graph shows median intersister (K-K) distance (bands show 2.5%, 50% and 97.5% quantiles) of all timely and lazy kinetochores throughout metaphase. **d**, Graph shows average autocorrelation (standard errors with 0.95 confidence interval) for the metaphase oscillations of all timely and lazy kinetochores. Negative depth of the autocorrelation curve indicates the regularity of kinetochore oscillations. **e**, Graphs show regression analyses of changes in the eight metaphase-anaphase variables (summary statistics) with respect to the maximum laziness throughout anaphase exhibited by lazy kinetochores pooled from cells treated with DMSO (grey, *n* = 14 lazy kinetochores), 2h noc (dark pink, *n* = 29 lazy kinetochores) or 4h noc (light pink, *n* = 48 lazy kinetochores). Black lines denote a linear fit to the data via maximum likelihood estimation, and the grey envelope shows the 95% confidence region for predictions from the model. *R*-squared and *p* values are shown for significantly correlating variables; for all variables see Supp. Table 1.

We reasoned that the intersister (K-K) distance may also scale with the severity of laziness. We found that among lazy kinetochores the reduction in K-K distance does indeed correlate with an increasing maximum laziness (*p* < 10^−3^, Fig. 4e). Furthermore, a decreasing oscillation amplitude and speed, and an increasing anaphase onset delay and twist also significantly correlate with increasing maximum laziness (Fig. 4e). Oscillation amplitude and speed are not significantly different when comparing the medians of timely vs. lazy kinetochores (Fig. 4a), which likely reflects the considerable heterogeneity in oscillatory dynamics (Fig. 1e). However, when restricted to only the lazy kinetochore population, correlations with maximum laziness are significant for these parameters, suggesting that increased laziness of a kinetochore is associated with impaired metaphase oscillations. These trends are consistent with the small number of lazy kinetochores (*n* = 22) in DMSO treated cells and therefore do not reflect any issues associated with the nocodazole treatment (Supp. Fig. 4a). Crucially, these data identify a dynamic metaphase signature which is associated with subsequent segregation fate during anaphase.

### Predicting kinetochore fate in anaphase based on its metaphase dynamics

Can this metaphase signature be used to successfully forecast whether a kinetochore will be lazy in the subsequent anaphase (Fig. 5a)? To address this question, we fitted a logistic regression model to determine which variables are most influential in predicting whether a kinetochore will be timely or lazy in anaphase, see Methods. Among covariates describing metaphase, the intersister (K-K) distance was the most influential variable because its model coefficient has the largest magnitude; and the 95% confidence interval [− 0.040 − 0.014] does not contain zero. Confidence intervals for other variables indicate weaker evidence for the amplitude, twist and the distance from the metaphase plate for an influence on the prediction (Fig. 5b). We trained this simple predictive model on the pooled dataset of cells treated with DMSO, 2h noc or 4h noc (*N* = 153 cells), and tested it on a separate set of DMSO-treated cells (*N* = 32), examining the predictive potential of single and multiple covariates (Fig. 5c-f). The area under the ROC curve is 0.65 for the full model on the test data (Fig. 5d,f) suggesting that the model clearly outperforms random chance. K-K distance is the most informative, performing to a similar extent to the full model, whilst the other metaphase variables do have some predictive capacity (Fig. 5d,f). Furthermore, K-K distance alone is a more powerful predictor on the test data than the timing of anaphase onset (Fig. 5d,f). Kinetochore dynamics in metaphase therefore enables forecasting of whether a kinetochore will be lazy during the subsequent anaphase.

**Figure 5.**
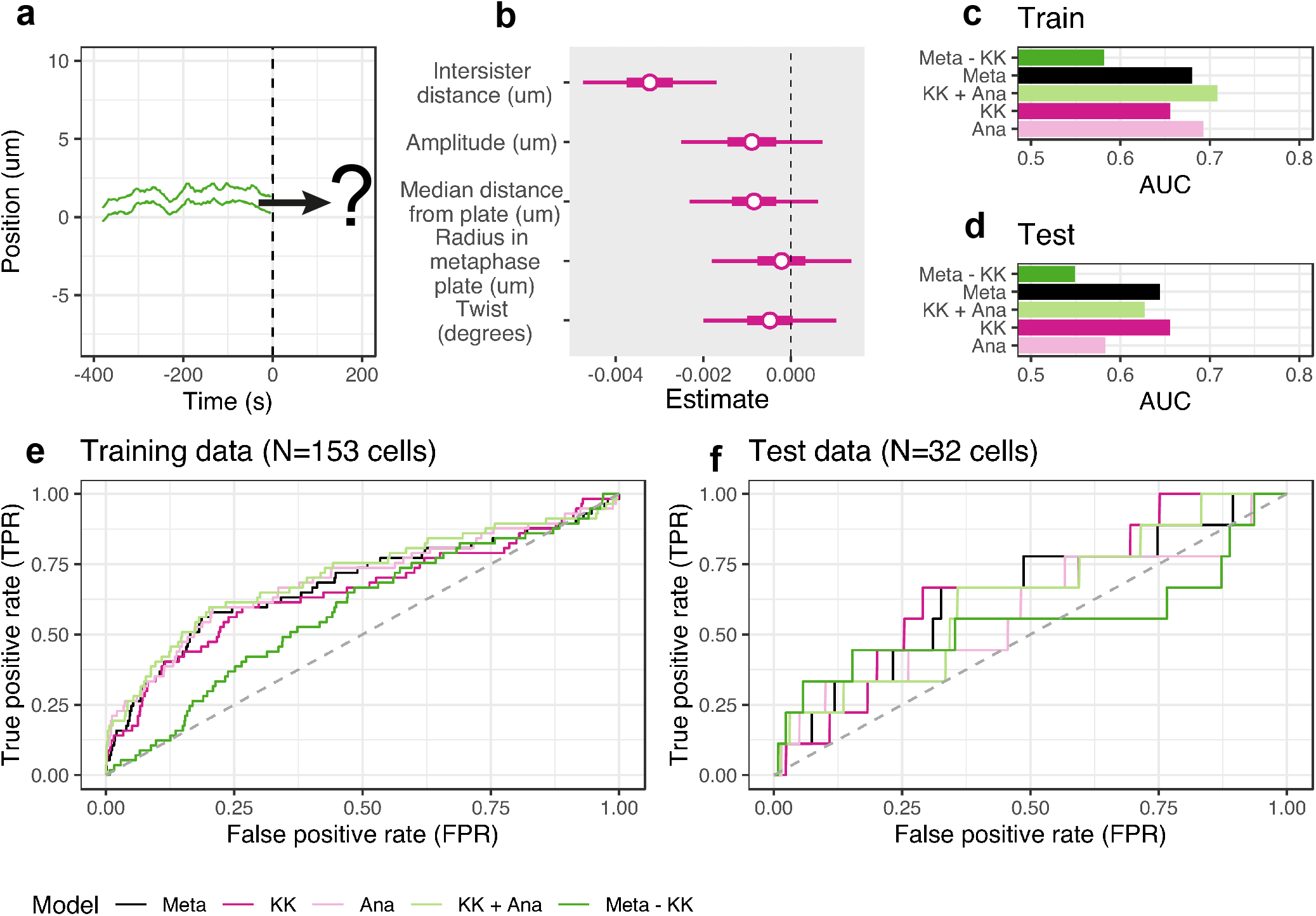
Predicting kinetochore fate in anaphase based on its metaphase dynamics. **a**, The metaphase trajectories of a sister pair (*x*-axis position in time) may forecast anaphase behavior of the two kinetochores. We consider model-based prediction of the fate of a kinetochore (lazy or timely) during segregation in anaphase based on the dynamics of the knetochore in metaphase. **b**, Plot shows estimated coefficients of metaphase covariates in the predictive model, as obtained by maximum likelihood estimation using pooled DMSO, 2h nocodazole, and 4h nocodazole treated cells. Coefficients indicate the influence of each covariate on the log-odds of whether a kinetochore is lazy. Dashed line indicates zero, and the influence of variables increases as the coefficient moves away from zero. **c**, Bar chart compares AUC (area under curve, the performance indicator that quantifies how good the performance of the predictive model is) for several models in their ability to classify lazy kinetochores on the training dataset (*N* = 153 cells; DMSO (*N* = 53 cells); 2h nocodazole (*N* = 46 cells); 4h nocodazole (*N* = 54 cells)). Meta: all metaphase variables shown in (b); KK: intersister (K-K) distance; Ana: anaphase onset time of a sister pair relative to the median anaphase onset for the cell; Meta - KK: all metaphase variables without K-K distance; Ana + KK: relative time of anaphase onset and K-K distance. **d**, Bar chart compares AUC of several models for their predictive capacity on the test dataset (DMSO; *N* = 32 cells); models as (c). **e**, ROC curves from which the AUCs given in (c) are obtained. **f**, ROC curves from which the AUCs given in (d) are obtained.

### Evidence for anaphase error correction process

Plotting the laziness for individual kinetochores over time shows how most of the kinetochores segregate in a timely fashion (Fig. 6a; grey areas). Moreover, a majority of the lazy kinetochores are seen to reduce their laziness over time (magenta trajectories), while high laziness persists for a minority of kinetochores (green trajectories). We hypothesize that a correction mechanism operates during anaphase to resolve lazy kinetochores. To define anaphase correction, we calculated whether the frame-to-frame laziness, *z*, of a kinetochore reduced below the laziness threshold, *a*, within 300 s of anaphase onset - approximately the timescale for the end of anaphase A^56,57^, (henceforth referred to as early anaphase). In DMSO treated cells, 93% of lazy kinetochores exhibit transient lazy behavior and are corrected in early anaphase (Fig. 6b and see example k1 in Fig. 6e, f). However, a small proportion of lazy kinetochores (7%) escape early anaphase correction, and persist beyond this time window; thus, these kinetochores are considered as uncorrected (for example see k1 and k2 in Fig. 6g, h). In nocodazole treated cells, the proportion of uncorrected kinetochores is 3.3 times higher (23% in 4h noc) and increases in a treatment duration dependent manner (Fig. 6a, b). After an initial increase in the number of lazy kinetochores following anaphase onset, the number of uncorrected lazy kinetochores decreases over time consistent with correction (Fig 6c).

**Figure 6.**
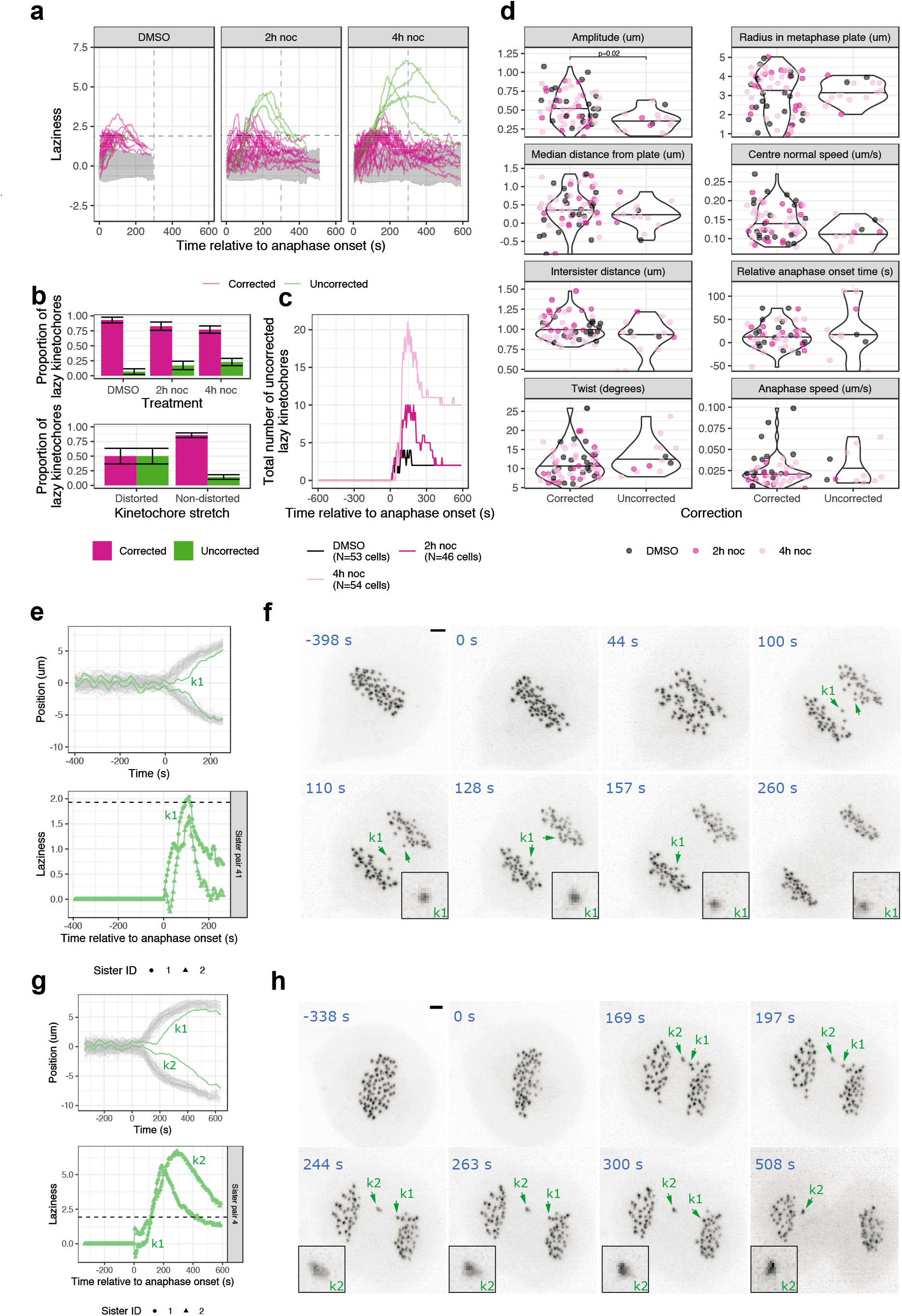
Correction of lazy chromosomes in anaphase is associated with dynamic kinetochore oscillations in metaphase. **a**, Graphs show laziness trajectories of lazy kinetochores (with > *a* maximum laziness) plotted over time. Black dashed line denotes laziness threshold (*a* = 1.93); grey dashed line indicates 300 s after anaphase onset (end of anaphase A). Grey area indicates 2.5% and 97.5% quantiles of laziness corresponding to trajectories of timely kinetochores (with < *a* maximum laziness). Magenta trajectories are the lazy kinetochores that are corrected (laziness decreased below < *a*) within 300 s of anaphase onset. Green trajectories are the lazy kinetochores that are not corrected (laziness not decreased below < *a*) within 300 s of anaphase onset. **b**, (top panel) Bar chart shows the proportions of corrected and uncorrected lazy kinetochores from cells treated with DMSO (*N* = 53 cells), 2h nocodazole (*N* = 46 cells) or 4h nocodazole (*N* = 54 cells). (bottom panel) Graph shows the proportions of corrected and uncorrected lazy kinetochores (*n* = 91; pooled from cells treated with DMSO, 2h nocodazole or 4h nocodazole) with distorted spots (n=14, 50% corrected) or non-distorted spots (*n* = 77, 86% corrected) during anaphase (*p* = 0.006). **c**, Bar chart shows total number of uncorrected lazy kinetochores remaining with laziness > *a* throughout anaphase. **d**, Violin plots show eight metaphase-anaphase variables (summary statistics) in lazy kinetochores (with > *a* maximum laziness) pooled from cells treated with DMSO, 2h nocodazole or 4h nocodazole, and classified as corrected (laziness decreased below < *a* within 300 s of anaphase onset) or uncorrected. Average values are median. **e**, (top panel) Track overlay (*x*-axis position in time) shows the trajectory of a transiently lazy kinetochore k1 (and its sister) that is corrected within 300 s of anaphase onset. (bottom panel) Graph shows the laziness trajectory of k1 kinetochore (and its sister) throughout anaphase. **f**, *Z*-projected movie stills of the cell in (e), where kinetochore k1 and its sister are annotated with green arrows. Zoomed images show that k1 is not distorted during anaphase. Scale bar is 2 µm. **g**, (top panel) Track overlay (*x*-axis position in time) shows the trajectory of a lazy kinetochore k2 (and its sister) that is not corrected. (bottom panel) Graph shows the laziness trajectory of k2 kinetochore (and its sister) throughout anaphase. **h**, *Z*-projected movie stills of the cell in (g), where kinetochore k2 and its sister are annotated with green arrows. Zoomed images show that k2 is distorted during anaphase, indicating merotelic attachment. Scale bar is 2 µm.

We reasoned that the corrected kinetochores may be different from the uncorrected population, potentially as a result of the microtubule-kinetochore attachment status that was established pre-anaphase. We thus compared the corrected and the uncorrected populations of lazy kinetochores and found that the latter have a significantly reduced amplitude of oscillation (25% average reduction, from 0.5 to 0.38 µm) in metaphase (Fig. 6d). Other variables do not show clear differences between the populations. This suggests that while a low K-K distance can predict subsequent lazy segregation, it is the kinetochores with damped oscillatory dynamics (reduced amplitude) that are more likely to avoid correction and fail to undergo accurate chromosome segregation. Examples of a lazy kinetochore with high amplitude oscillations that is subsequently corrected and a lazy kinetochore with impaired oscillations that fails to be corrected are displayed in Fig 6e, f (Movie S3) and Fig 6g, h (Movie S4) respectively.

In addition, we assessed whether lazy kinetochores are distorted during anaphase. Previous studies showed that a merotelic kinetochore can undergo distortion (stretching) due to the correctly and incorrectly attached microtubules pulling it towards opposite poles^15,16^. Our high temporal resolution imaging (4.7s/frame) enables us to capture kinetochore stretching and recoiling events, that manifest as the distortion of kinetochore spot shape. In fact, manual assessment of 3D movies showed that 14 of the lazy kinetochores (*n* = 91) underwent distortion at varying degrees. Because of the opposing forces acting on merotelically attached kinetochores, we use spot distortion as evidence for merotely. We assessed whether the distorted lazy kinetochores differentiate from the non-distorted lazy kinetochores in terms of their ability to be corrected during anaphase. The distorted population (e.g. Fig. 6g, h) of lazy kinetochores are less likely to be corrected, 50% versus 86% (e.g. non-distorted lazy kinetochore; Fig. 6e, f) (Fig. 6b (bottom panel)). This indicates that merotelic attachment is associated with a failure of error correction during early anaphase.

### Aurora B inhibition disrupts the correction of chromosome segregation errors during anaphase

Because Aurora B kinase activity is required for pre-anaphase error correction ^30^, we tested whether it is involved in the early anaphase error correction process outlined above. To do this, we used the small molecule inhibitor ZM447439 of Aurora kinase (ZM)^33^ following washout from nocodazole or DMSO. We only imaged the cells that were exposed to the Aurora inhibitor after chromosomes had completed congression at the spindle equator; in other words, after they have “passed” Aurora B dependent error correction in prometaphase. Inhibition of Aurora B in this way still allowed anaphase to initiate, but led to an increase in the number of lazy kinetochores (Fig. 7a,b). We note that Aurora B inhibition does slow the overall separation of kinetochore clusters (Fig. 7c), which is consistent with the anaphase roles of Aurora B reported previously^58,59^.

**Figure 7.**
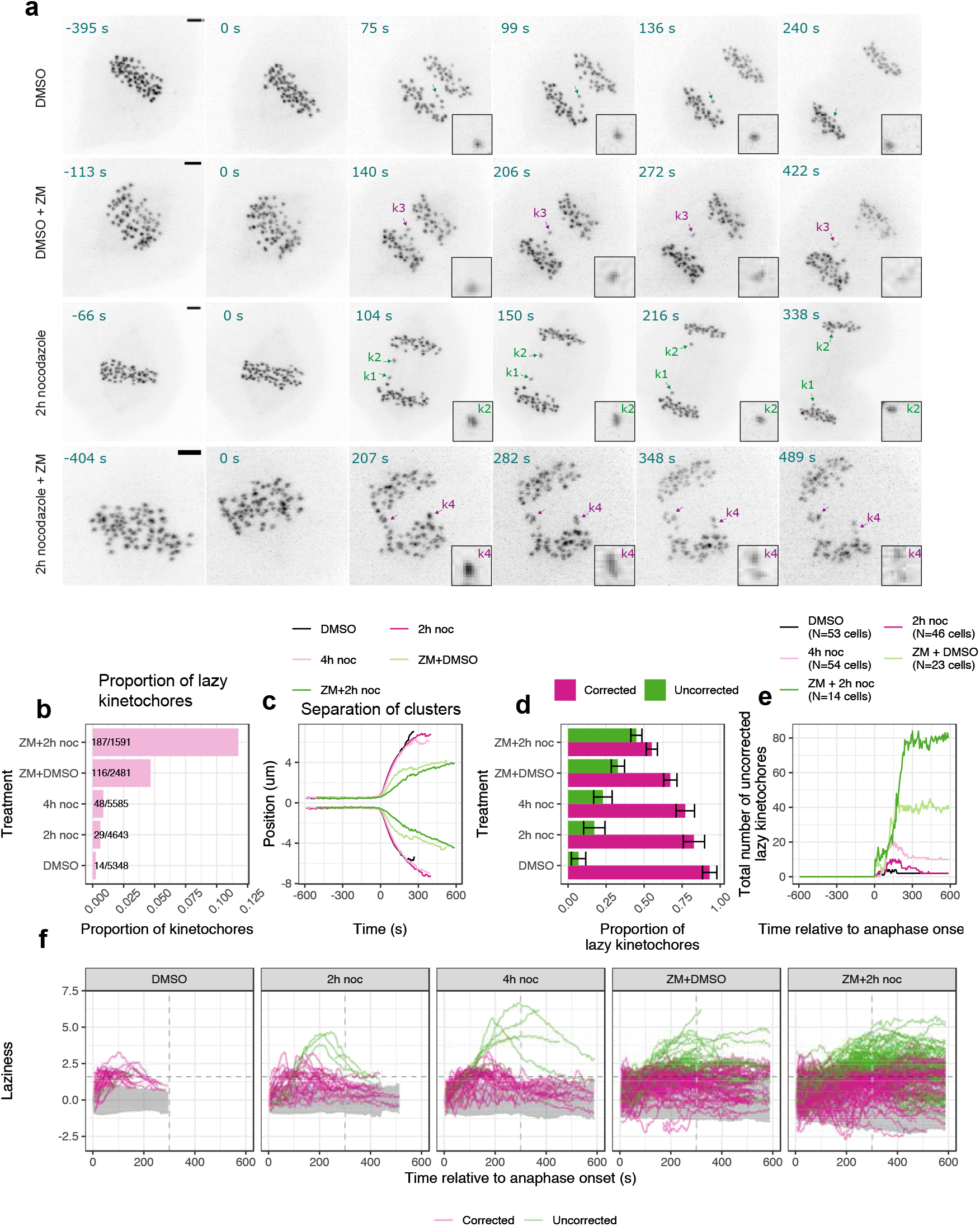
Aurora B kinase activity is required for correcting lazy kinetochores during anaphase. **a**, *Z*-projected movie stills of a representative (top panel) cell that was treated with DMSO washout, which had multiple transient lazy kinetochores (annotated with green arrow) that are corrected in early anaphase; (second panel from top) cell that was treated with DMSO washout prior to being exposed to Aurora inhibitor (ZM), which had a lazy kinetochore k3 (annotated with purple arrow) that was distorted and split into two parts, but was not corrected in anaphase; (third panel from top) cell that was treated with 2h nocodazole washout, which had lazy kinetochores k1 and k2 (annotated with green arrows), that were transiently distorted (and recoiled) and corrected in early anaphase; (bottom panel) cell that was treated with 2h nocodazole washout prior to being exposed to Aurora inhibitor (ZM), which had multiple lazy kinetochores (k4 annotated with purple arrow, distorted and split into two parts) and not corrected in anaphase. Scale bar is 2 µm. **b**, Bar chart shows proportions (and numbers) of lazy kinetochores from different treatment groups. **c**, Track overlay (*x*-axis position in time) shows the average position of segregating kinetochore clusters from all cells in a treatment group to compare the speed of kinetochore cluster segregation. **d**, Bar chart shows the proportions of corrected and uncorrected lazy kinetochores (with > *a* maximum laziness) from cells treated with DMSO (*N* = 53 cells); 2h noc (*N* = 46 cells); 4h noc (N=54 cells); DMSO and then ZM (*N* = 23 cells); 2h noc and then ZM (*N* = 14 cells). **e**, Graph shows total number of uncorrected lazy kinetochores remaining with laziness > *a* throughout anaphase. Lazy kinetochores with laziness that fails to fall below the threshold, *a* by the end of the movie are assumed to remain uncorrected. **f**, Laziness trajectories of lazy kinetochores (with > *a* maximum laziness) plotted over time. Black dashed line denotes laziness threshold (*a* = 1.93); grey dashed line indicates 300 s after anaphase onset. Grey area indicates 2.5% and 97.5% quantiles of laziness corresponding to trajectories of timely kinetochores (with < *a* maximum laziness). Magenta trajectories annotate the lazy kinetochores that are corrected (laziness decreased below *a*) within 300 s of anaphase onset. Green trajectories annotate the lazy kinetochores that are not corrected (laziness does not decrease below *a*) within 300 s of anaphase onset.

Loss of Aurora B activity dramatically disrupted the correction of lazy kinetochores in anaphase (Fig. 7d, e). For example, Fig. 7a shows two lazy kinetochores (*k*3 and *k*4) that failed to be corrected, displaying severe spot distortion. In fact, these kinetochores later disintegrated due to persistent pulling forces towards opposite poles. This is in contrast, to cells with active Aurora B in which most lazy kinetochores are corrected during early anaphase (Fig. 7d; see example kinetochores *k*1 and *k*2 in Fig. 7a which are corrected after transient spot distortion). Importantly, this effect was also observed in the cells that were treated with DMSO prior to Aurora B inhibition, demonstrating that the lack of anaphase error correction is not a consequence of nocodazole washout. There is a synergistic effect of nocodazole washout and Aurora B inhibition elevating the total number of lazy kinetochores (Fig. 7b) and the fraction that were uncorrected (Fig. 7d).

This failure to correct lazy kinetochores under Aurora B inhibition is also evident when plotting the laziness trajectories for individual kinetochores over time (Fig. 7e, f). In cells with active Aurora B, even in those cases where laziness fails to fall below threshold by 300 s (classed as uncorrected) the correction of lazy kinetochores appears to be in progress (green trajectories are declining, Fig. 7f). In contrast, under Aurora inhibition (ZM), lazy kinetochores have a laziness that typically does not have a clear decline phase and remains high until the end of the movie (green trajectories show no obvious decline, Fig. 7f). These differences, and the dynamics of early anaphase correction when Aurora B is active, are clearly visible in plots of the total number of uncorrected kinetochores over time (Fig. 7e). These data suggest that Aurora B activity is required for correcting chromosome segregation errors in early stages of anaphase. This further supports the idea that many more kinetochores than previously thought are at risk of mis-segregation, and that cells are reliant on anaphase error correction processes to avoid aneuploidy.

## Discussion

The scoring of the “lagging chromosome” phenotype underpins hundreds of cell biology studies on the genes and processes that give rise to chromosome mis-segregation. The lagging chromosome event has also gained widespread use as a proxy for the occurrence of merotelic kinetochore-microtubule attachments. Despite this, there is no convention on what constitutes a lagging chromosome. Most studies rely on fixed cell imaging which can only provide a snapshot of anaphase and is therefore sensitive to the “moment” in anaphase and unavoidable observer bias i.e. when does a chromosome become lagging? This is particularly imprecise when using chromosome labels, since the position of bulky chromosome arms do not always represent the position of kinetochores where microtubules attach to segregate them. Recent efforts to quantify lagging chromosomes from fixed cell analyses are an important step forward^60^. However, these approaches do not use a time-series, and therefore cannot capture the temporal and spatial evolution of short-lived events. Here, we quantify dynamic behaviors of kinetochores as they progress through metaphase and anaphase. To set apart our dynamic approach from the studies based on fixed cell imaging we have chosen to avoid the term “lagging”, and instead propose the use of “laziness”; a novel measure that quantifies the segregation behavior of an individual kinetochore through time.

This analysis reveals that there is a substantial population of kinetochores that become lazy during anaphase. 18% of DMSO treated cells possess at least one kinetochore that became lazy. This is considerably higher than the proportions of cells with lagging chromosomes (5-7%) in previous reports based on fixed cell imaging^22,24,28^. Crucially, we found that 93% of these lazy kinetochores were corrected in anaphase A, in part explaining the underestimation by fixed cell imaging. Our algorithm uses a threshold, tuned to a false positive rate of ∼5% against manual assessment. This threshold gives a ∼30% false negative rate which implies we are underestimating the lazy population; this is predominantly because our algorithm is not able to detect lazy kinetochores with low levels of Ndc80 binding (low signal-to-noise ratio) nor does it include some highly stretched (merotelic) kinetochores that are close to the cluster (insufficiently lazy). Remarkably, this suggests that mal-orientated kinetochores are not a rare event but rather a common feature of a normal unperturbed mitosis. Anaphase error correction mechanisms (this study and^16^) and “checkpoints” that delay nuclear envelope reassembly^47,61^ are therefore crucial for cells to minimize the probability of lazy chromosomes persisting into telophase where they may form micronuclei and undergo further mutational processes^6^ (see below for discussion).

Our data show that a lowered K-K distance (a proxy for tension across the centromeric chromatin) and perturbed regularity of oscillations in metaphase are a signature for lazy behavior of kinetochores during anaphase. Moreover, the severity of laziness correlates with the extent of K-K distance reduction along with lower oscillation speed and amplitude. Crucially, our predictive model is able to forecast lazy kinetochores in a new dataset of unperturbed cells. Thus, our findings are not restricted to cells that are arrested and released from nocodazole (the majority of the training data). Together, this indicates that problems in chromosome segregation are already set before anaphase onset.

What does the metaphase signature represent at the molecular level? Our data do not support a role for precocious sister chromatid separation (PSCS) because this would be expected to increase K-K distances, and is normally the result of longer duration mitotic arrests^24^. This also suggests that our nocodazole-arrest-and-release procedure is unlikely to affect cohesion between sister chromatids. The most natural interpretation of our data is that the metaphase signature reflects dynamic behavior of merotelically attached kinetochores in metaphase. Two observations would support this: 1) it is well established that nocodazole arrest-and-release increases the number of lagging chromosomes with merotelic attachments^13^, and our experiments also show a similar increase in both the number of lazy kinetochores and in the level of their laziness, 2) merotelic attachment of a kinetochore would be consistent with a reduction in its K-K distance due to the pulling forces from the incorrect attachment, bringing the merotelic kinetochore closer to its sister during metaphase. However, we observed only 15% (14 in 91) of lazy kinetochores with qualitatively detectable physical distortion (stretching) in the Ndc80 signal – an established feature of merotely^36^. The metaphase signature for lazy kinetochores may therefore also reflect other attachment configurations, or underlying defects (i.e. chromosome entanglement) that can give rise to lazy segregation.

At this stage we propose the following working model: during metaphase sister kinetochores are expected to bind up to 20 microtubules from opposite spindle poles (Fig. 8, i). A considerable proportion of these kinetochores would, however, have a merotelic configuration with reduced K-K distance and reduced distance from the metaphase plate. The severity of merotely would range from one or two mis-attached microtubules on the incorrect side (mero-amphitelic), through to full occupancy with equivalent numbers on the incorrect and correct sides of a kinetochore (balanced-merotelic; Fig. 8, ii)^14^. It is tempting to speculate that the number of mis-attached microtubules correlates with the laziness in anaphase. We found that the most severe laziness associates with disrupted metaphase oscillations, which would be consistent with these merotelic kinetochores having balanced attachment to the two spindle poles. As the cell progresses into anaphase, kinetochores with few mis-attached microtubules (mero-amphitelic) would be rapidly corrected (Fig. 8, step iii to iv); while those with more severe merotely (balanced-merotelic) would be distorted (stretched) and possibly persist to telophase (Fig. 8, step v to vi). This would also be consistent with previous microtubule poison based experiments that led to a model in which reduced kinetochore-microtubule occupancy (in metaphase) is associated with reduced K-K distance and an increase in lagging chromosomes^62^. The idea is that with fewer microtubules on the correct side there is an increased chance of balanced merotely.

**Figure 8.**
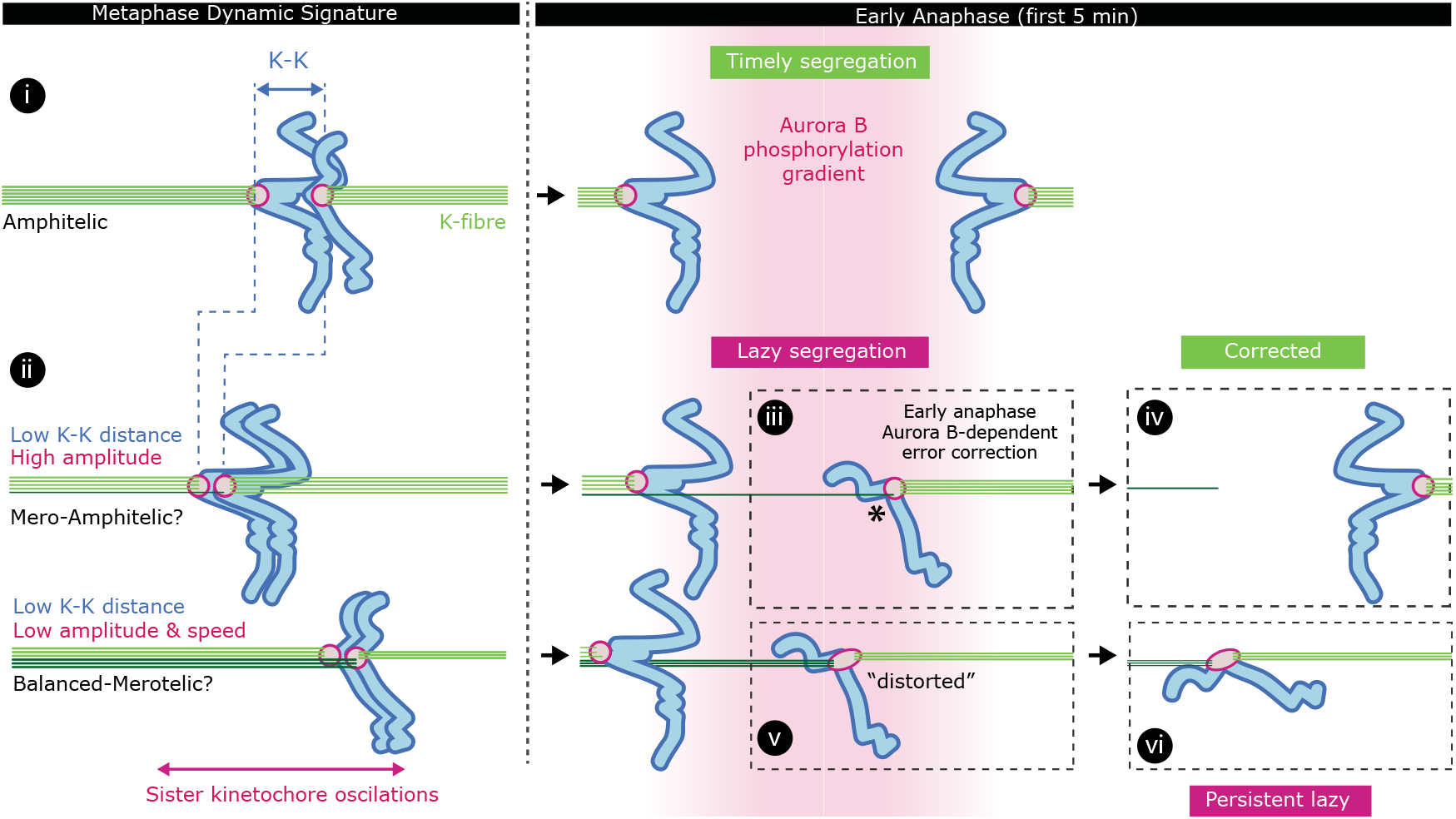
Kinetochore life histories reveal the origins of chromosome mis-segregation and correction mechanisms. Working model explaining how attachment status and metaphase dynamics of kinetochores can affect their segregation behavior in anaphase. In metaphase, most kinetochores have amphitelic attachments (step i). However, a considerable proportion of kinetochores can have merotelic configuration ranging in severity from mero-amphitelic, with fewer mis-attached microtubules, to balanced-merotelic, with equal numbers of microtubules on the correct and incorrect sides of a kinetochore (step ii). In anaphase, mero-amphitelic kinetochores can be rapidly corrected via the Aurora B phosphorylation gradient (step iii and iv); whereas balanced-merotelic kinetochores become distorted (stretched) and possibly persist to telophase (step v and vi).

Our data reveal that Aurora B activity is involved in the active correction of lazy kinetochores in early anaphase. This is distinct from any proposed roles of Aurora B in late anaphase^63,64^, and defines a new layer of error correction. In pre-anaphase cells, Aurora B located within the centromere-kinetochore mediates the destabilization of improper microtubule-kinetochore attachments, which are normally under low tension, rendering the kinetochores unattached^65^. However, Aurora B predominantly relocates to the spindle midzone in anaphase, where it generates a phospho-gradient. After separating from its sister at anaphase onset, the incorrect attachment side of a merotelic lazy kinetochore is more likely to be closer to the midzone (asterisk in Fig. 8, step iii), hence within the Aurora B phospho-gradient. Thus, Aurora B may destabilize the microtubules on the incorrect side more efficiently. If both sides of a merotelic lazy kinetochore are within the midzone phospho-gradient, Aurora B may reduce the stability of all microtubule attachments on both sides equally, and catalyze the error correction through a tug-of-war between correct and incorrect sides, leading to the ultimate removal of incorrect attachment. This model is consistent with our observation that balanced merotelics (stretching) are less likely to correct (50% correction), while mero-amphitelics are efficiently corrected (86% correction) in early anaphase. In this regard, the probability of correction is also forecast by the behavior of a lazy kinetochore in metaphase – most notably by high oscillation amplitude (distance travelled along the spindle axis). We suggest that higher amplitudes reflect sister kinetochores with fewer mis-attached microtubules (mero-amphitelic), which is consistent with findings reported in^16^. These kinetochores would be more efficiently corrected by Aurora B (Fig. 8, Step iii). We would also predict that mero-syntelic attachments (with fewer attachments to the correct pole than to the incorrect pole) would “correct”, but result in non-disjunction due to earlier removal of the thinner correct attachment^29^. An important next step will be to establish dynamic signatures for these merotelic attachment variations, and to track their fate and origin during mitosis.

In conclusion, we have established a quantitative framework to define kinetochore segregation behaviors, and have dissected the mechanisms that cause and correct lazy kinetochores in anaphase. This has revealed how kinetochores’ behavior in metaphase forecasts their future, and that a high proportion of kinetochores are at risk of mis-segregation. We have defined a new layer of error correction, which operates in anaphase to limit chromosome mis-segregation. This work provides firm ground for further investigations into the origins of whole chromosome aneuploidies – a hallmark of tumorigenesis and reproductive failure in humans.

## Methods

### Cell culture, drug treatment and generation of cell lines

Immortalized (hTERT) diploid human retinal pigment epithelial (RPE1) cell line (MC191), expressing endogenously tagged Ndc80-eGFP, was generated by CRISPR-Cas9 gene editing^49^. hTERT-RPE1 cells were grown in DMEM/F-12 medium containing 10% fetal bovine serum (FBS), 2 mM L-glutamine, 100 U/ml penicillin and 100 mg/ml streptomycin (full growth medium); and were maintained at 37°C with 5% CO_2_ in a humidified incubator. For nocodazole arrest-and-release experiments cells were treated with DMSO (1/50,000 v/v) or 330 nM nocodazole for 2h or 4h, followed by transferring the coverslip to 5 ml full growth medium without any drugs, and 20 min incubation (washout step). After the washout, the coverslip was transferred to the lattice light sheet microscope (LLSM) bath filled with CO_2_-independent L15 medium, where live imaging takes place. For Aurora B kinase inhibitor (ZM447439) treatment cells were treated with DMSO (1/50,000 v/v) or 330 nM nocodazole for 2h, followed by transferring the coverslip to 5 ml full growth medium without any drugs, and 30 min incubation (washout step). After the washout, coverslip was transferred to the LLSM bath filled with CO_2_-independent L15 medium, including 4 µM ZM447439, where live imaging takes place.

### Live imaging by Lattice light sheet microscope (LLSM)

The lattice light sheet microscope^48^ used in this study was manufactured by 3i (https://www.intelligent-imaging.com). Cells were seeded on 5 mm radius glass coverslips one day before imaging. On the imaging day, cells were treated with drugs, and the coverslip was transferred to the LLSM bath filled with CO2-independent L15 medium, where live imaging takes place. The LLSM light path was aligned at the beginning of every imaging session by performing beam alignment, dye alignment and bead alignment, followed by the acquisition of a bead image (at 488 nm channel) for measuring the experimental point spread function (PSF). This PSF image is later used for the deconvolution of images. 3D time-lapse images (movies) of Ndc80-eGFP were acquired at 488nm channel using 1% laser power, 50 ms exposure time/z-plane, 93 z-planes, 307 nm z-step, which results in 4.7 s/z-stack time(frame). Acquired movies were de-skewed and cropped in XYZ and time, using Slidebook software in order to reduce the file size. Cropped movies were then saved as OME-TIFF files in ImageJ.

### Manual assessment of lazy kinetochores

Movies for each cell were manually assessed by searching for prominent lagging kinetochores using the 3D view mode of Slidebook software (to avoid projection effects in the *Z* direction). Cells that had rotated on the coverslip during image acquisition were reoriented such that their segregation axes would correspond to the direction perpendicular to the metaphase plate according to the observer. This ensures an optimal cell orientation for viewing the segregation dynamics, and thus was used for the manual assessment. A list of cells ranked by maximum laziness (detected in each cell) was compared (see Supp. Fig. 3a) with the results of manual inspection of the same data (*N* = 153 cells; DMSO, 2h noc, 4h noc pooled). This revealed that the algorithm identified kinetochores with high laziness (threshold = ∼2 based on extremal analysis) within the subpopulation of cells that had been manually recorded as having zero prominent lagging kinetochores. Manual reassessment of these cells revealed that 12 of them did indeed have lagging kinetochores which had not been noticed in the first manual inspection. Cells in the ranked list were then reassessed until it was clear no further lagging kinetochores were found. Any additional cells with lagging kinetochores were included in the population of cells classified as having manually assigned lagging kinetochores. This list was used to calibrate the algorithm’s laziness threshold.

### Deconvolution and kinetochore tracking

Movies were deconvolved using the Richardson-Lucy algorithm for deconvolution via the Flowdec library^66^. The PSF used was a non-isotropic 3D Gaussian PSF fitted to the measured experimental PSF from each imaging session. Gaussian noise similar to background was added to blank regions of the image to avoid artefacts at the boundaries to blank image regions. Kinetochore tracking (KiT v2.3) software^50^ was used to detect and track kinetochores, and subsequently pair sister kinetochores. Detection is achieved via adaptive thresholding of movies, and refined via a Gaussian mixture model. Detected kinetochores are linked between frames to form tracks via a Kalman filter and linear assignment problem. Tracks are grouped based on metaphase dynamics via a linear assignment problem. A plane is fitted to the metaphase plate as a reference coordinate system, in which the *x* direction points perpendicular to the metaphase plate, and *y* and *z* lie within the plate.

### Mechanistic anaphase model

A hierarchical model was used to describe the positions of each kinetochore pair in a cell. The model takes the form of a stochastic differential equation with terms for the spring force due to chromatin connecting sister kinetochores, the polar ejection force, and forces due to microtubule polymerization/depolymerization, as in previous work^67^. In metaphase, the following dynamics hold for the position, 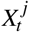, of sister kinetochore *j* at time *t*:

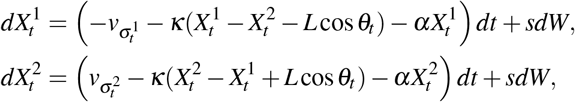

which comprises the mechanical forces 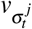 due to K-fibre polymerisation/depolymerisation associated with the hidden K-fibre state 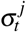 (polymerising (+), depolymerising (-)), a centromeric spring force between the sisters, 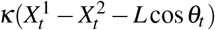, spring constant *κ* and natural length *L*, projected to the *x*-axis (sister-sister twist *θ*_*t*_), the polar ejection forces with proportionality constant, *α*, and the thermal fluctuations with standard deviation *s*.

In anaphase, the polar ejection force and the chromatin spring force are assumed to be absent, giving:

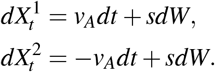

with anaphase speed *ν* _*A*_. An additional anaphase reversal state, only accessible from the anaphase state, is included in the model to account for reversals in anaphase such that

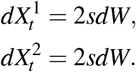

where we assume kinetochores are diffusing.

The K-fibre polymerisation state is described by 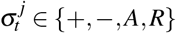, in metaphase, either polymerising (+) or depolymerising (-), and in anaphase, either moving towards their pole (*A*) or in a reversal reversal (*R*). This discrete hidden state evolves as a Markov process. Biophysical parameters are assumed individual to each kinetochore pair, while parameters governing the transitions between hidden states are assumed to be shared between all kinetochore pairs in a cell giving a hierarchical cell based model.

To fit the hierarchical anaphase model to experimental data, we take a Bayesian approach and draw samples from the posterior distribution via Markov chain Monte Carlo (MCMC). Specifically, we use the No-U-Turn-Sampler (NUTS)^68^ implementation of Hamiltonian Monte Carlo^69^ via the software Stan^70^. The likelihood is evaluated via the forward algorithm^71^. Convergence of MCMC chains is assessed via the Gelman-Rubin 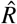 statistic^72,73^ using only results where 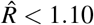 1.10 for all parameters. Across all treatment groups, MCMC chains converged for 224/259 cells, and among these cells for 7040/7253 kinetochore pairs (Supp. Fig. 1 d,e) which were used in subsequent analysis.

The mechanistic anaphase model was fitted to long trajectories from each cell, annotating the trajectory by sister direction and anaphase separation time for each pair. Based on the estimates of anaphase onset times, we obtained an estimate of the median time of anaphase onset for a cell. Using this estimate of the median time of anaphase onset for a cell, we assessed the laziness for all tracked kinetochores (including kinetochore pairs with short tracks, and unpaired kinetochores).

### Definition of the laziness, *z*

The laziness of a kinetochore at time, *t*, is given by

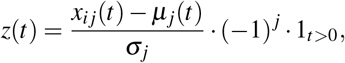

where *x*_*i j*_(*t*) is the position of kinetochore *j* from sister pair *i* (measured relative to the metaphase plate), *µ*_*j*_(*t*) is the median position of the daughter cell cluster *j*, and *σ*_*j*_ is a scale for the spread of daughter cell cluster *j* estimated via the median absolute deviation on 20 early anaphase frames. The 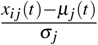 term is similar to the definition of a *z* score with reference to a normal distribution. The (−1) ^*j*^ term ensures that *z*(*t*) is positive for kinetochores between clusters, i.e. indicative of slower segregation than the median. The indicator term 1_*t*>0_ makes *z*(*t*) equal to 0 prior to anaphase onset (at *t* ≤ 0).

### Summary statistics to describe dynamics of kinetochores in metaphase and anaphase

The intersister (K-K) distance (see Fig. 1c) is calculated in 3D for a kinetochore pair as 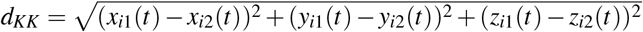 where *x*_*i j*_(*t*), *y*_*i j*_(*t*), *z*_*i j*_(*t*) are the position in each coordinate of kinetochore *j* from sister pair *i* at time *t*, with the *x* coordinate perpendicular to the metaphase plate. All metaphase summary statistics are calculated across a trajectory excluding the 60 s prior to anaphase onset, and summarised via the median. To calculate the amplitude of oscillation of an individual kinetochore (see Fig. 1c), we used a sliding window of 20 frames and calculated the amplitude as *A* = (max(*x*_*i j*_(*t*)) − min(*x*_*i j*_(*t*))) */*2. The average distance from the metaphase plate is calculated as *d*_*MP*_ = (− 1)^1+ *j*^median(*x*_*i j*_(*t*)) using a signed distance to indicate perpendicular distance from the metaphase plate in the direction of the cluster towards which the kinetochore will segregate. The radius within the metaphase plate is calculated as 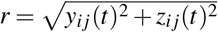 The centre normal speed of a kinetochore pair is calculated as the framewise speed of the mean position of the pair as follows *ν*_*CNS*_ = (*x*_*i j*_(*t* + Δ*t*) + *x*_*i j*_(*t* + Δ*t*)) − (*x*_*i*1_(*t*) + *x*_*i*2_(*t*))*/*2Δ*t* where Δ*t* is the time step between frames, and this is summarised via the standard deviation across a trajectory to capture the scale of changes in speed over an oscillation. The twist angle of a kinetochore pair is computed as cos^−1^(median(|cos(*ϕ*)|)), where cos(*ϕ*) = (*x*_*i*2_(*t*) − *x*_*i*1_(*t*))*/* ∥ **x**_*i*2_(*t*) − **x**_*i*1_(*t*) ∥ with **x**_*ij*_(*t*) = (*x*_*i j*_(*t*), *y*_*i j*_(*t*), *z*_*ij*_(*t*))^*T*^. The relative anaphase onset time onset is calculated as 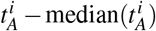 −median(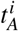) where the median is calculated across kinetochore pairs, and 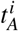 is the median estimate of anaphase onset for a pair based on the mechanistic anaphase model. The anaphase speed, *ν*_*A*_, is a 1D speed in the direction perpendicular to the metaphase plate, as in the mechanistic anaphase model described above.

### Logistic regression model

The logistic regression model is a generalized linear model to express the relationship between a binary dependent variable, *y*, (here corresponding to whether a given kinetochore has laziness above the threshold, *a* = 1.93) and a matrix, **X**, of covariates (summary statistics describing dynamics of the given kinetochore in metaphase only). Suppose that *p* = *P*(kinetochore is lazy). We assume a linear relationship between the predictor variables and the log-odds, 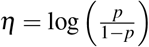, of a kinetochore being lazy such that *η* = **b**_**0**_ + **b**^*T*^ **X**. We obtain

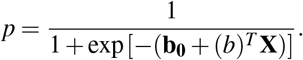

We consider five models based on different combinations of covariates: 1) Metaphase covariates (K-K distance, amplitude, median distance from metaphase plate, radius in metaphase plate, twist); 2) K-K distance only; 3) time of anaphase onset relative to the median for a cell, only; 4) K-K distance and time of anaphase onset relative to the median for a cell; 5) Metaphase covariates without K-K distance (amplitude, median distance from metaphase plate, radius in metaphase plate, twist). Each model is fitted based on maximum likelihood estimation to data from *N* = 153 cells (*n* = 10160 kinetochores; DMSO, 2h noc, 4h noc pooled). Predictions are made for all kinetochores in *N* = 32 DMSO cells unseen by the models to evaluate performance.

### Statistical comparisons

Differences in medians were assessed via two sample Wilcoxon tests. Differences in proportions were assessed via Fisher’s exact test. Correction for multiple testing was performed via the Holm-Bonferroni method. All tests were performed using the rstatix package in the software R v3.5.2.

## Supporting information

3D view movie of the cell shown in Fig. 1.

Z-projected (maximum intensity) movie of the cell shown in Fig. 2d, in which k1 kinetochore and its sister are annotated.

Z-projected (maximum intensity) movie of the cell shown in Fig. 6f, in which k1 kinetochore and its sister are annotated.

Z-projected (maximum intensity) movie of the cell shown in Fig. 6h, in which k2 kinetochore and its sister are annotated.

## Acknowledgements

We thank Sarah McClelland for comments on the manuscript and Helena Coker from the Computational and Microscopy Development Unit (CAMDU) for support with lattice light sheet microscopy. The Lattice Light Sheet Microscope Facility was established at Warwick with a Wellcome Trust Multi-user Equipment grant to AM (grant 208384/Z/17/Z). OS, JH, NB and AM are supported by BBSRC (BB/R009503/1). AM is also supported by a Wellcome Trust Senior Investigator Award (grant 106151/Z/14/Z).

## Author contributions statement (CRediT)

Project was conceived by AM, NJB. OS designed and performed all experiments, lattice light-sheet (LLS) imaging and image preparation. JUH devised and wrote the analysis software, developing KiT for LLS tracking and the trajectory annotation metaphase-anaphase model. OS, JUH analysed the data. Manuscript written by OS, JUH, AM, NJB.

## Supplementary Material

**Movie S1**. 3D view movie of the cell shown in Fig. 1.

**Movie S2**. Z-projected (maximum intensity) movie of the cell shown in Fig. 2D, in which k1 kinetochore and its sister are annotated.

**Movie S3**. Z-projected (maximum intensity) movie of the cell shown in Fig. 6F, in which k1 kinetochore and its sister are annotated.

**Movie S4**. Z-projected (maximum intensity) movie of the cell shown in Fig. 6H, in which k2 kinetochore and its sister are annotated.

**Table 1.** Supplementary Table 1

| sumstat | adj.r.squared | p.value |
| --- | --- | --- |
| Amplitude (um) | 0.07 | 0.0095 |
| Intersister distance (um) | 0.15 | <0.001 |
| Centre normal speed (um/s) | 0.11 | 0.0015 |
| Anaphase speed (um/s) | 0.02 | 0.15 |
| Median distance from plate (um) | -0.01 | 0.76 |
| Radius in metaphase plate (um) | -0.01 | 0.6 |
| Twist (degrees) | 0.08 | 0.0073 |
| Relative anaphase onset time (s) | 0.32 | <0.001 |

**Figure S1.**
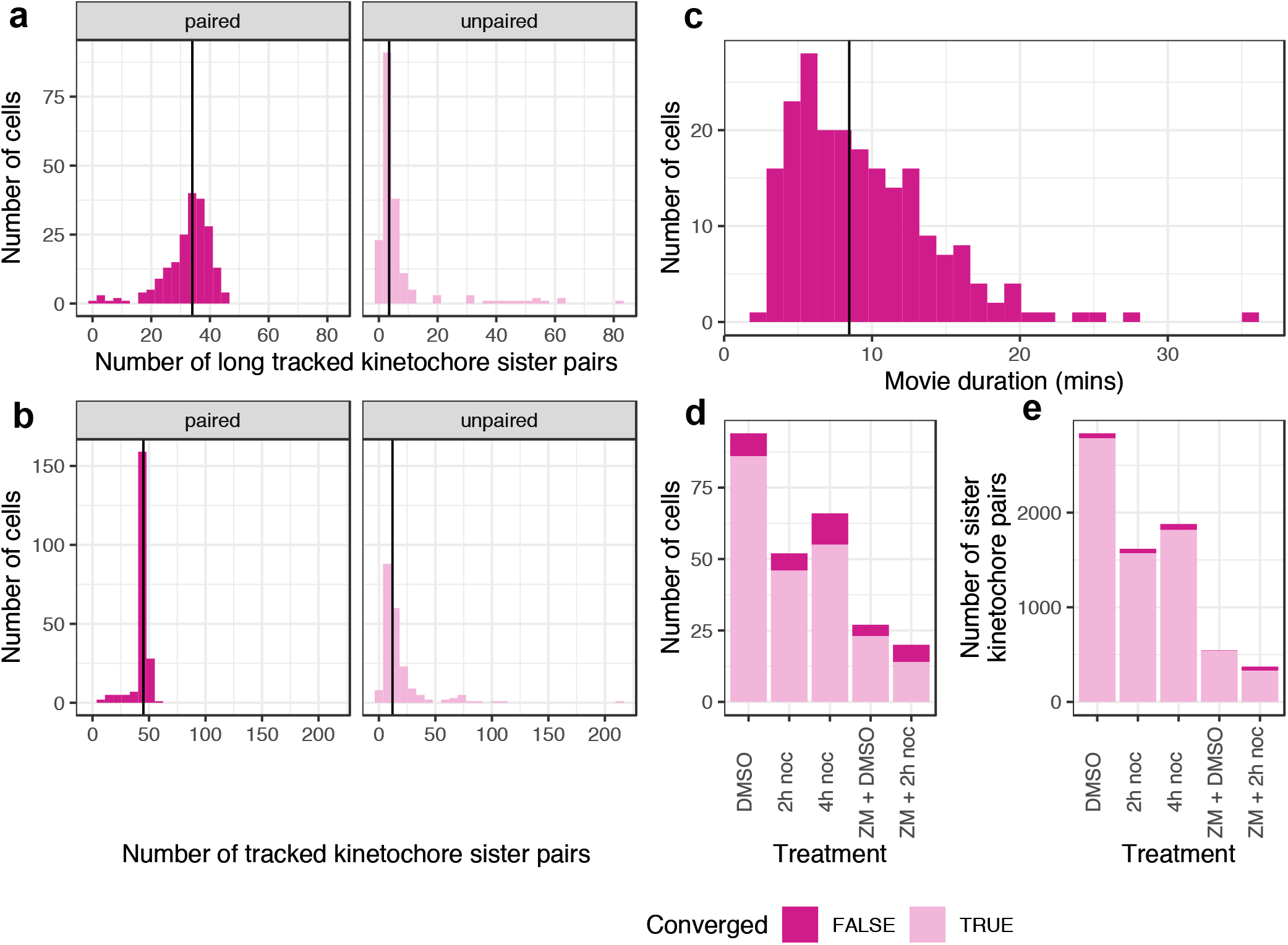
Near complete tracking of kinetochore sister pairs over long timescales. **a**, Histograms show the number of cells, in which long tracks (>75% of the 3D time series (movie) for each cell) were obtained for the given number of kinetochore sister pairs (paired) or single kinetochores (unpaired). **b**, Histograms show the number of cells, in which tracks (irrespective of their length in a movie) were obtained for the given number of kinetochore sister pairs (paired) or single kinetochores (unpaired). **c**, Histogram shows the number of cells with the given movie duration. Black lines indicate median in (a), (b) and (c). **d**, Bar chart shows the number of cells from each treatment group for which MCMC chains have converged for the mechanistic anaphase model. **e**, Bar chart shows the number of pairs (within cells that have converged) from each treatment group for which MCMC chains have converged for the mechanistic anaphase model.

**Figure S2.**
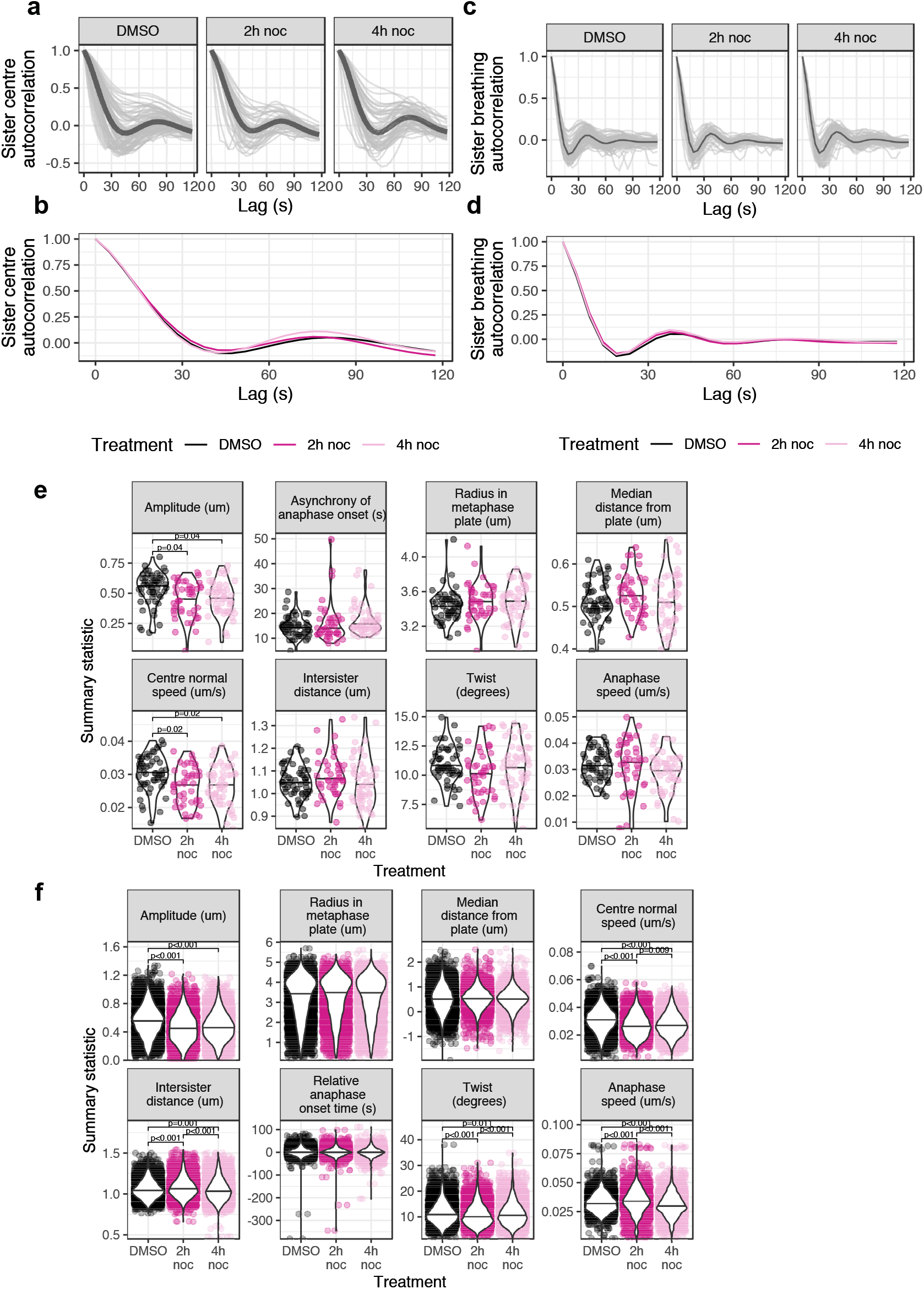
Nocodazole arrest-and-release has little impact on metaphase oscillation dynamics, except for a reduction in oscillation amplitude. **a**, Graphs compare the sister centre (mean position of sister kinetochores) autocorrelation for kinetochore oscillations in treatment groups. Each of the light grey curves represents the average (of all kinetochores in that cell) autocorrelation for a cell. Dark grey curve indicates the average autocorrelation for kinetochore oscillations in all cells per treatment group. **b**, Graph shows the average sister centre autocorrelations for the treatment groups plotted together. **c**, Graphs compares the sister breathing autocorrelation for kinetochore pairs in treatment groups, which is a readout for the regularity of breathing movement between sister kinetochores during their oscillations. Each of the light grey curves represents the average (of all kinetochores in that cell) autocorrelation for a cell. Dark grey curve indicates the average autocorrelation for kinetochore breathing in all cells per treatment group. **d**, Graph shows the average sister breathing autocorrelations for the treatment groups plotted together. **e**, Graphs compare eight metaphase-anaphase variables (summary statistics) in the cells (average of kinetochores per cell) treated with DMSO (*N* = 53 cells), 2h nocodazole (*N* = 46 cells) or 4h nocodazole (*N* = 54 cells). Average values are median. **f**, Graphs compare eight metaphase-anaphase variables (summary statistics) in kinetochores from the cells treated with DMSO (*n* = 5348 kinetochores), 2h nocodazole (*n* = 4643 kinetochores) or 4h nocodazole (*n* = 5585 kinetochores). Average values are median.

**Figure S3.**
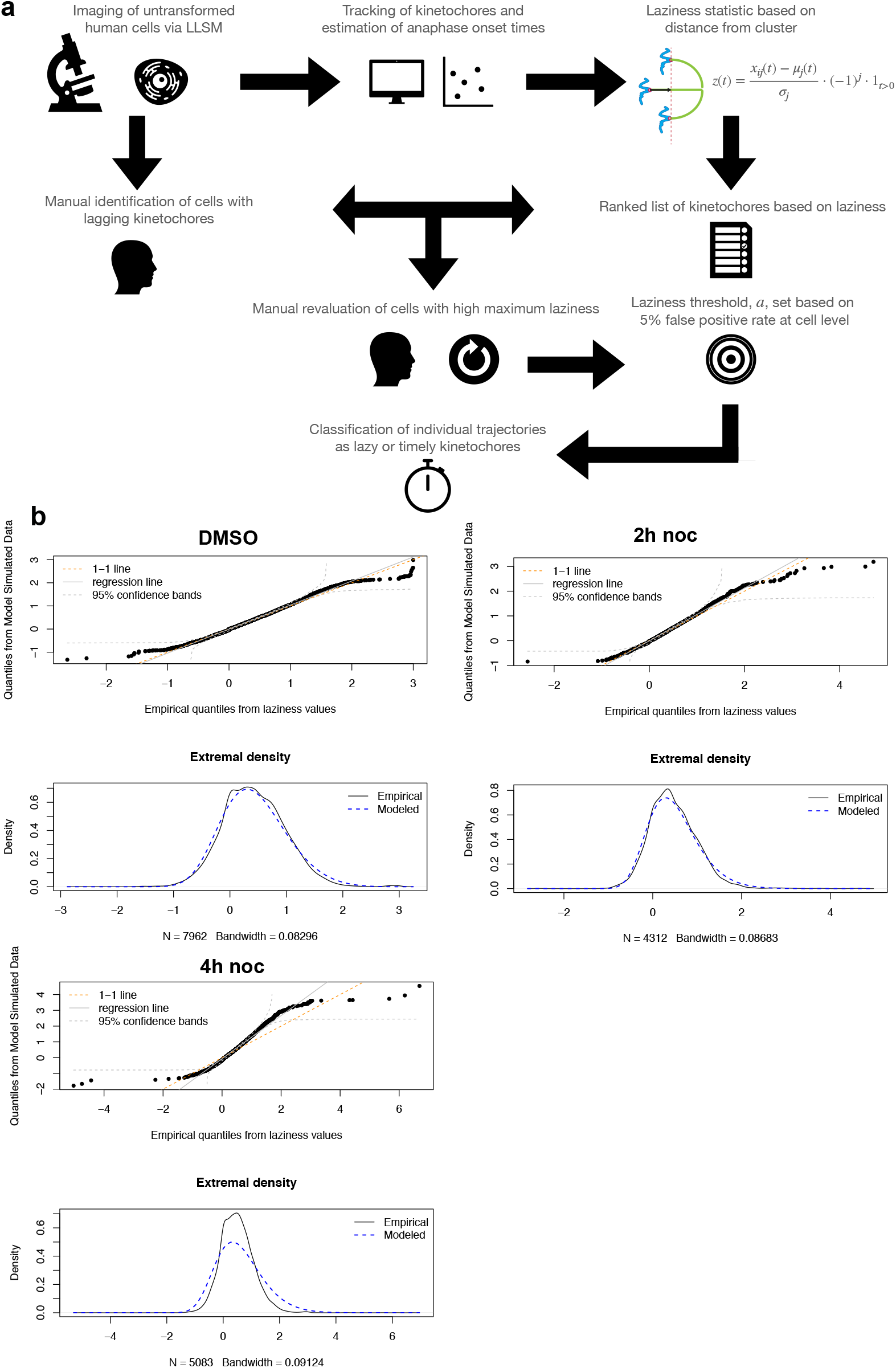
Estimation of laziness threshold and its calibration to manual assessment. **a**, Schematic illustrates the data analysis pipeline: data collection (LLSM imaging), tracking of kinetochores and calculation of laziness for each of them, calibration of laziness threshold in the light of manual assessments and automated analyses, and classification of individual kinetochores based on their segregation behavior. **b**, Quantile-quantile (Q-Q) plots show the maximum laziness scores fitted to a generalized extremal value distribution, which suggest that beyond a laziness threshold, a, of approximately 2, the quality of the fit breaks down for kinetochores from the cells treated with DMSO (*n* = 5348 kinetochores), 2h nocodazole (*n* = 4643 kinetochores) or 4h nocodazole (*n* = 5585 kinetochores).

**Figure S4.**
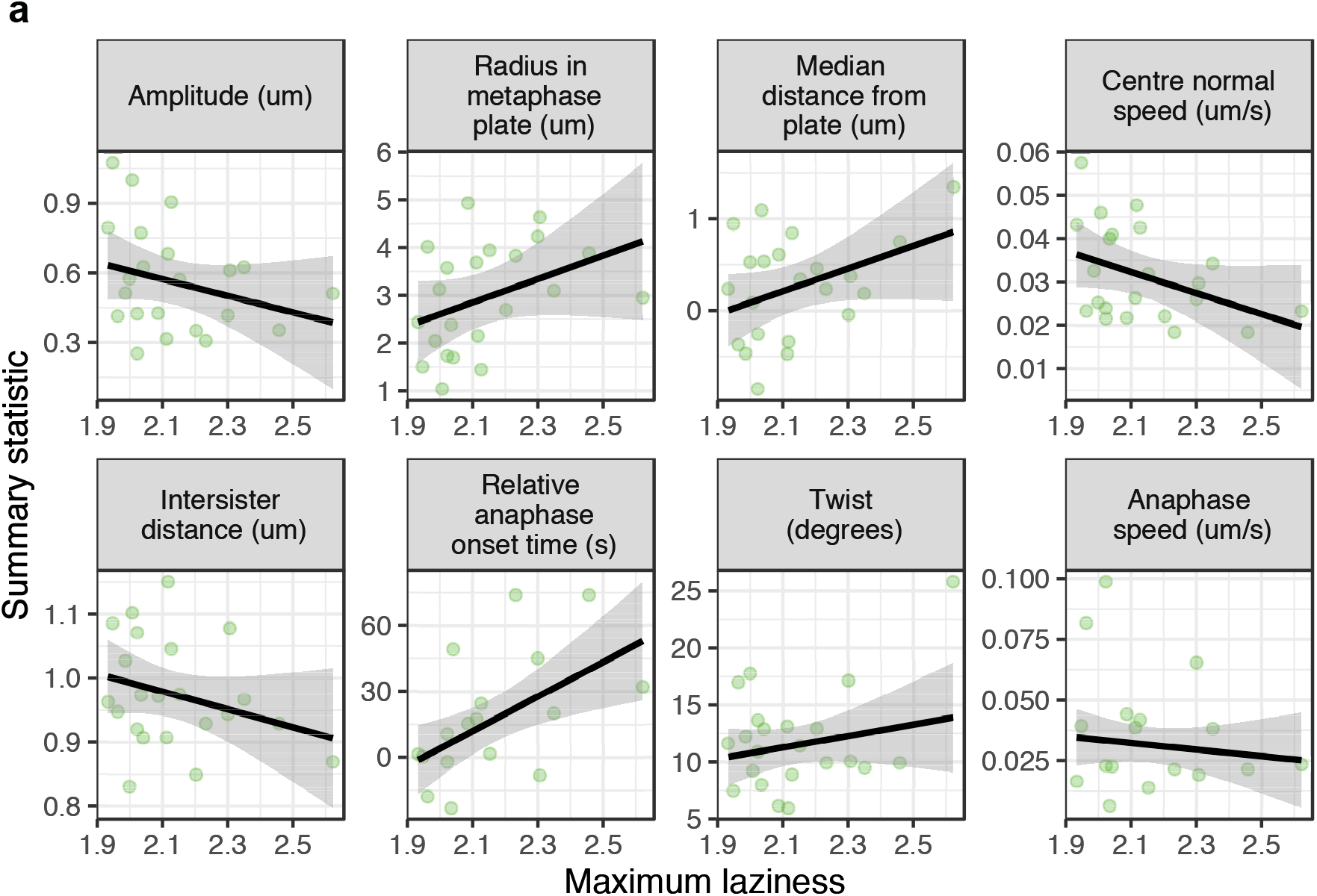
Metaphase signature is consistent with the lazy kinetochores from unperturbed cells. **a**, Graphs show regression analyses of changes in the eight metaphase-anaphase variables (summary statistics) with respect to the maximum laziness throughout anaphase exhibited by lazy kinetochores from the cells treated with DMSO (*n* = 22 lazy kinetochores). Black lines denote a linear fit to the data obtained via maximum likelihood estimation and the grey envelopes show the 95% confidence interval for predictions.

**Table 2.**
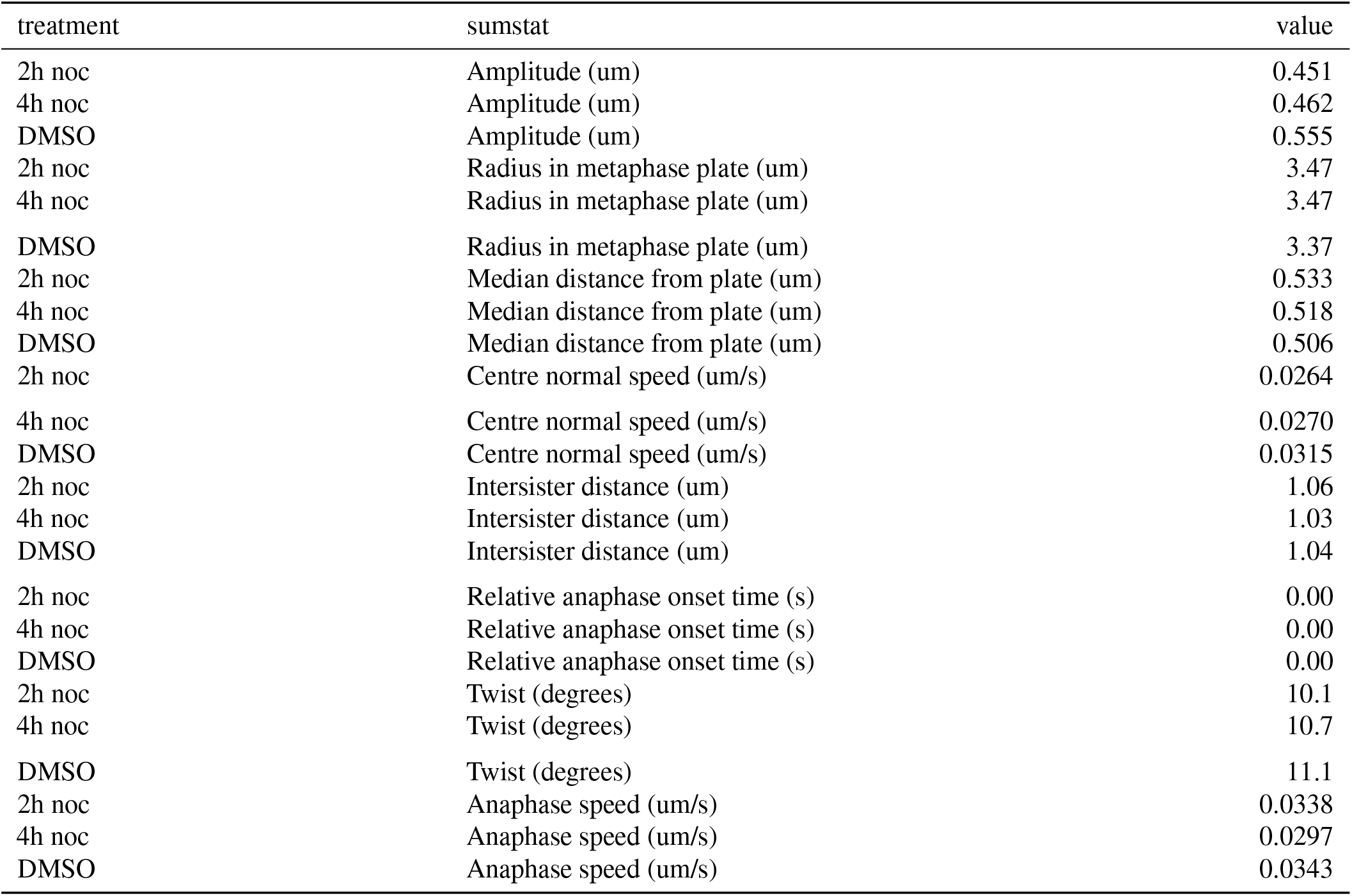
Supplementary Table 2

| treatment | sumstat | value |
| --- | --- | --- |
| 2h noc | Amplitude (um) | 0.451 |
| 4h noc | Amplitude (um) | 0.462 |
| DMSO | Amplitude (um) | 0.555 |
| 2h noc | Radius in metaphase plate (um) | 3.47 |
| 4h noc | Radius in metaphase plate (um) | 3.47 |
| DMSO | Radius in metaphase plate (um) | 3.37 |
| 2h noc | Median distance from plate (um) | 0.533 |
| 4h noc | Median distance from plate (um) | 0.518 |
| DMSO | Median distance from plate (um) | 0.506 |
| 2h noc | Centre normal speed (um/s) | 0.0264 |
| 4h noc | Centre normal speed (um/s) | 0.0270 |
| DMSO | Centre normal speed (um/s) | 0.0315 |
| 2h noc | Intersister distance (um) | 1.06 |
| 4h noc | Intersister distance (um) | 1.03 |
| DMSO | Intersister distance (um) | 1.04 |
| 2h noc | Relative anaphase onset time (s) | 0.00 |
| 4h noc | Relative anaphase onset time (s) | 0.00 |
| DMSO | Relative anaphase onset time (s) | 0.00 |
| 2h noc | Twist (degrees) | 10.1 |
| 4h noc | Twist (degrees) | 10.7 |
| DMSO | Twist (degrees) | 11.1 |
| 2h noc | Anaphase speed (um/s) | 0.0338 |
| 4h noc | Anaphase speed (um/s) | 0.0297 |
| DMSO | Anaphase speed (um/s) | 0.0343 |

